# A new variance ratio metric to detect the timescale of compensatory dynamics

**DOI:** 10.1101/742510

**Authors:** Lei Zhao, Shaopeng Wang, Lauren M. Hallett, Andrew Rypel, Lawrence W. Sheppard, Max C.N. Castorani, Lauren G. Shoemaker, Kathryn L. Cottingham, Katharine Suding, Daniel C. Reuman

**Affiliations:** Beijing Key Laboratory of Biodiversity and Organic Farming, College of Resources and Environmental Sciences, China Agricultural University, Beijing 100193, China; Department of Ecology and Evolutionary Biology and Kansas Biological Survey, University of Kansas, Higuchi Hall, 2101 Constant Ave, Lawrence, KS 66047, USA; Department of Ecology, College of Urban and Environmental Science, and Key Laboratory for Earth Surface Processes of the Ministry of Education, Peking University, Beijing 100080, China; Environmental Studies Program and Department of Biology, University of Oregon, Eugene, OR 97403, USA; Department of Wildlife, Fish, & Conservation Biology, University of California Davis, Greensboro, NC 27402, USA; Department of Environmental Sciences, University of Virginia, Charlottesville, VA 22904, USA; Botany Department, University of Wyoming, Laramie, WY, 82071, USA; Department of Biological Sciences, Dartmouth College, Hanover, NH 03755, USA; Department of Ecology & Evolution Biology, University of Colorado Boulder, CO, 80303, USA; Laboratory of Populations, Rockefeller University, 1230 York Ave, New York, NY, 10065, USA

**Keywords:** variance ratio, timescale, community stability, compensatory dynamics, synchrony, tsvr

## Abstract

Understanding the mechanisms governing ecological stability – why a property such as primary productivity is stable in some communities and variable in others – has long been a focus of ecology. Compensatory dynamics, in which anti-synchronous fluctuations between populations buffer against fluctuations at the community level, is a key theoretical mechanism of stability. Classically, compensatory dynamics have been quantified using a “variance ratio” approach that compares the ratio between community variance and aggregate population variance, such that a lower ratio indicates compensation and a higher ratio indicates synchrony among species fluctuations. However, population dynamics may be influenced by different drivers that operate on different timescales, and evidence from aquatic systems indicates that communities can be compensatory on some timescales and synchronous on others. The variance ratio and related metrics cannot reflect this timescale-specificity, yet have remained popular, especially in terrestrial systems. Here, we develop a timescale-specific variance ratio approach that formally decomposes the classical variance ratio according to the timescales of distinct contributions. The approach is implemented in a new R package, called tsvr, that accompanies this paper. We apply our approach to a long-term, multi-site grassland community dataset. Our approach demonstrates that the degree of compensation versus synchrony in community dynamics can vary by timescale. Across sites, population variability was typically greater over longer compared to shorter timescales. At some sites, minimal timescale-specificity in compensatory dynamics translated this pattern of population variability into a similar pattern of greater community variability on longer compared to shorter timescales. But at other sites, differentially stronger compensatory dynamics at longer compared to shorter timescales produced lower-than-expected community variability on longer timescales. Within every site there were plots that exhibited shifts in the strength of compensation between timescales. Our results highlight that compensatory versus synchronous dynamics are intrinsically timescale-dependent concepts, and our timescale-specific variance ratio provides a metric to quantify timescale-specificity and relate it back to the classic variance ratio.

## Introduction

The stability of ecosystem functions is central to the reliable provisioning of ecosystem services (Oliver *et al.*, 2015), and understanding mechanisms underlying ecological stability is a fundamental question in ecology (MacArthur, 1955). A key insight into ecosystem dynamics is that stable aggregate ecosystem functions such as total productivity can be composed of highly variable component parts (Gonzalez and Loreau, 2009). For example, compensatory dynamics stabilize productivity when different populations have offsetting fluctuations, such that increases in the abundance or biomass of one or more species are accompanied by corresponding decreases in others (e.g., Schindler, 1990, Frost *et al.*, 1995, Bai *et al.*, 2004, Hallett *et al.*, 2014). Conversely, when species increase or decrease together (e.g., when species share responses to an external driver), the resulting synchrony increases aggregate community variability and has been used as a proxy for instability (e.g., Houlahan *et al.*, 2007, Keitt, 2008, Feio *et al.*, 2015, Ma *et al.*, 2017).

Population fluctuations are ubiquitous. Consequently, characterizing patterns of species fluctuations over time and in relation to each other is essential to understand stability. Importantly, population fluctuations can be shaped by a variety of drivers that operate on different timescales. For example, acute or pulsed disturbances are a strong predictor of population dynamics in some systems (e.g., heavy rainfall events or cold winter events regulating insect dynamics; Mutshinda *et al.*, 2011). In other systems, population dynamics are strongly driven by long-term climate cycles (Mantua *et al.*, 1997, Sheppard *et al.*, 2016). As a result, communities may exhibit a timescale specificity such that synchrony (or compensation) among species can occur at some timescales but not others (Keitt, 2008, Downing *et al.*, 2008). Moreover, the co-occurrence of short- and long-timescale drivers in the same system may result in communities that are synchronous over some timescales but compensatory over others (as idealized in Fig. 1).

**Figure 1.**
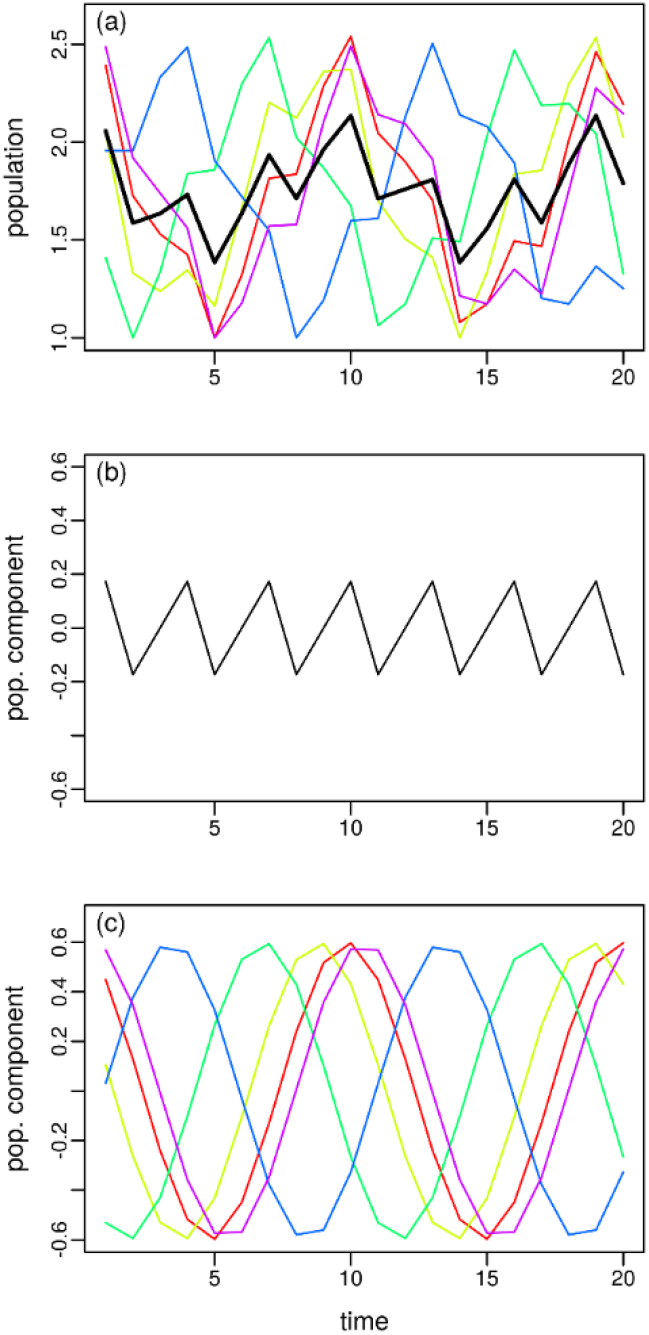
Introduction to concepts using an idealized example. Five population time series (panel a, colors distinguish the time series, the black line is the average) were constructed by summing a sine wave of period (timescale) 3 years and amplitude 0.2 (panel b) with five randomly and independently phase-shifted sine waves of period 10 years, amplitude 0.6 (panel c), and then vertically shifting each time series to have minimum value 1. Thus, the time series of panel a were constructed to be synchronous on the 3-year timescale and asynchronous (randomly related) on the 10-year timescale. However, the classical variance ratio (Peterson, 1975, Schluter, 1984) was 1.06, suggesting that the species show neither compensation nor synchrony across the entire time series, missing the deliberately constructed timescale-specific processes.

Evaluating timescale specificity in community variability has a tradition in aquatic ecology (Vasseur *et al.*, 2005, Keitt and Fischer, 2006, Vasseur and Gaedke, 2007, Downing *et al.*, 2008, Keitt, 2008, Vasseur *et al.*, 2014, Brown *et al.*, 2016), but methods used in that literature have typically relied on data-hungry techniques that may not be appropriate for shorter time series. For example, Vasseur *et al.* (2005) found that phytoplankton in Lake Constance showed compensatory dynamics at a sub-annual timescales, driven by grazing pressure and competition for nutrients during summer and fall, but synchronous dynamics at most other timescales. Downing *et al.* (2008) described how zooplankton in experimental ponds had synchronous dynamics at quite short timescales (~10-day periods), but had compensatory dynamics at longer timescales (~80-day periods). Overall, synchrony has tended to be more common than compensatory dynamics in freshwater plankton communities, and the timescales at which compensatory dynamics occur appear to be system-specific (Vasseur *et al.*, 2014, Brown *et al.*, 2016). Timescale-dependent approaches have facilitated a deeper understanding of community stability in freshwaters.

Compensatory dynamics have been the subject of much study and debate in terrestrial systems. Some studies find general evidence for compensatory dynamics (Bai *et al.*, 2004, Hector *et al.*, 2010), others find them to be context-dependent (Grman *et al.*, 2010, Hallett *et al.*, 2014, Xu *et al.*, 2015), and others conclude that synchronous dynamics dominate terrestrial systems (Houlahan *et al.*, 2007, Valone and Barber, 2008). To date, terrestrial ecologists have relied exclusively on metrics that do not incorporate timescale (most commonly, those developed by Schluter, 1984, Loreau and de Mazancourt, 2008, Hallett *et al.*, 2014). These types of analysis originated with the classic variance ratio, which can be calculated for the shorter time series available in these systems (Schluter, 1984). The variance ratio compares the variance of the aggregate community to the expected variance under the assumption of independent population fluctuations across the entire time series. As such, a variance ratio greater than one indicates that populations are generally synchronous, whereas a variance ratio less than one indicates compensation (Peterson, 1975, Schluter, 1984). This simplicity and ease of interpretation has held wide appeal across diverse scientific fields (e.g., Gotelli, 2000, Klug *et al.*, 2000, Houlahan *et al.*, 2007, Winfree and Kremen, 2009). However, the variance ratio cannot disentangle the timescales at which dynamics occur (Fig. 1), ultimately hindering possibilities for considering mechanisms of community dynamics.

Here we develop and apply a new timescale-specific variance ratio appropriate for terrestrial grasslands and other systems with shorter, regularly-spaced time series. In contrast to some of the timescale-specific approaches previously used on plankton data, our techniques provide a formal decomposition by timescale of the classic variance ratio approach, so that appropriately averaging/summing the components of the new approach across timescales recovers the classic non-timescale-specific results; one can then quantify the contributions of timescale bands as well as individual timescales. We first develop the theory that underlies our approach and provides timescale-specific measures of community and population variability. Second, we apply this theory to long-term (11–30 years) records of plant community composition at six grasslands across the United States (Hallett *et al.*, 2014). We address the fundamental questions of 1) whether synchrony/compensatory dynamics are timescale-dependent phenomena, and 2) whether compensatory dynamics are rare, compared to synchrony, in grassland systems, as appears to be the case for plankton systems (Vasseur *et al.*, 2005, Keitt and Fischer, 2006, Vasseur and Gaedke, 2007, Downing *et al.*, 2008, Keitt, 2008, Vasseur *et al.*, 2014, Brown *et al.*, 2016). We aim to demonstrate our timescale-specific approach in a way that deepens our understanding of grassland community dynamics as developed using the traditional variance ratio metric (Hallett *et al.*, 2014). Finally, we provide a software package for the R programming language to facilitate wider adoption of our techniques.

## Theory

Community dynamics are commonly described empirically using abundance (e.g., count, density, percent cover, or biomass) time series *x_i_*(*t*) for times *t* = 1,…*T* for the taxa *i* = 1,…,*S* comprising a community (e.g., all plant species in a quadrat). We denote *μ_i_* = mean (*x_i_*(*t*)) and *v_ij_* = cov (*x_i_*(*t*), *x_j_*(*t*)) as the means and covariances, respectively, of population dynamics through time. We denote *x*_tot_ (*t*) = Σ_*i*_*x_i_*(*t*) as the total population density or biomass time series, and *μ*_tot_ = Σ_*i*_*μ*_i_ and *v*_tot_ = Σ_*i,j*_*v_ij_* as the mean and variance, respectively, through time of *x*_tot_ (*t*).

We use the square of the coefficient of variation of 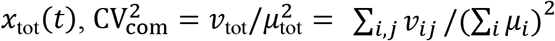, to quantify the dynamical variability of the summed community property (Wang and Loreau, 2014). If populations were entirely independent, the covariances *v_ij_* for *i* ≠ *j* would be 0 and 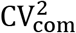 would equal 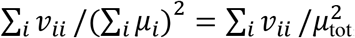, which we denote as 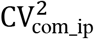. We refer to 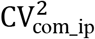 as the aggregate population dynamical variability of the system because it represents the degree of overall community variability that would result solely from the dynamics of individual taxa, discounting interactions among taxa.

The classic *variance ratio* 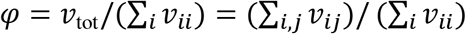 (Schluter, 1984) is well known to quantify the extent to which population fluctuations of different taxa reinforce each other (through synchrony of temporal fluctuations, *φ* > 1) or cancel each other (through compensation, *φ* < 1) in the community total time series *x*_tot_(*t*). The variance ratio satisfies 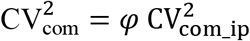 (theorem 2 of Appendix S2; Peterson, 1975, Schluter, 1984). Thus values *φ* > 1 (respectively, < 1) correspond to greater (resp., lesser) community variability than would be expected if dynamics of different taxa were independent, reflecting synchronous (resp., compensatory) dynamics.

Next, we develop timescale-specific statistics using spectral methods, which are a standard statistical tool in ecology (Reuman *et al.*, 2006, Vasseur and Gaedke, 2007, Defriez *et al.*, 2016) and many other fields. All definitions and computational choices about the basics of power spectra and cospectra are detailed in Appendix S1 in the Supporting Information. The power spectrum of the time series *x_i_*(*t*), here denoted *s_ii_*(*σ*) and defined for timescales of oscillation *σ* = *T*/(*T* – 1), *T*/(*T* – 2),…,*T*/2, *T* decomposes var(*x_i_*(*t*)) by timescale in the sense that *s_ii_*(*σ*) will be tend to be larger for timescales *σ* on which *x_i_*(*t*) is oscillating strongly. Thus, the power spectrum provides information on the dominant timescales of oscillation in *x_i_*(*t*). The power spectrum is a formal decomposition of variance across timescales because 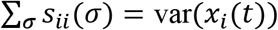 (Appendix S1). The cospectrum *s_ij_*(*σ*) of the time series *x_i_*(*t*) and *x_j_*(*t*) decomposes cov (*x_i_*(*t*), (*t*)) by timescales, i.e., *s_ij_*(*σ*) is defined for *σ* = *T*/(*T* – 1), *T*/(*T* – 2),…, *T*/2,…, *T* such that 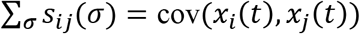 (Appendix S1), and *s_ij,_* (*σ*) tends to be larger for timescales on which the two time series predominately covary (i.e., they are varying synchronously, with substantial and largely in-phase periodic components at those timescales).

Community variability 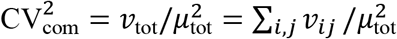 is made timescale-specific by replacing the covariances in the numerator with their timescale decompositions based on power spectra and cospectra, 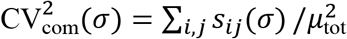. Thus 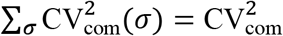 (theorem 1 in Appendix S2), and 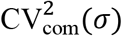 reveals to what extent fluctuations on each timescale contribute to community variability through time. Aggregate population variability 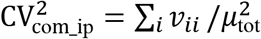 is made timescale-specific again by replacing the variances in the numerator with their timescale decompositions, 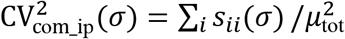. Thus 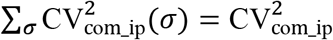 (theorem 1 in Appendix S2) and 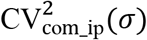 reveals to what extent fluctuations on each timescale contribute to aggregate population variability.

A timescale-specific variance ratio can be defined by replacing covariances in the definition of by their timescale-specific decompositions, 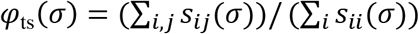. Thus the timescale-specific variance ratio satisfies 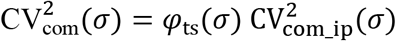. Values *φ*_ts_ (*σ*) > 1 correspond to synchrony at timescale *σ* and values *φ*_ts_(*σ*) < 1 correspond to compensatory dynamics at *σ*. We then define the quantity 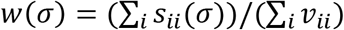, which represents the relative amount of variation in populations across timescales. Because of the normalization in the denominator of *w*(*σ*), Σ_σ_*w*(*σ*) = 1. We cannot simply sum *φ*_ts_ (*σ*) across timescales to recover*σ*. However, Σ_*σ*_*w* (*σ*) *φ*_ts_ (*σ*) = *φ*, so *φ* is instead a weighted average across timescales of *φ*_ts_ (*σ*) (Appendix S2). The relationships between the classic quantities 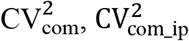 and *φ* and the timescale-specific extensions we have defined are summarized schematically in Fig. 2.

**Figure 2.**
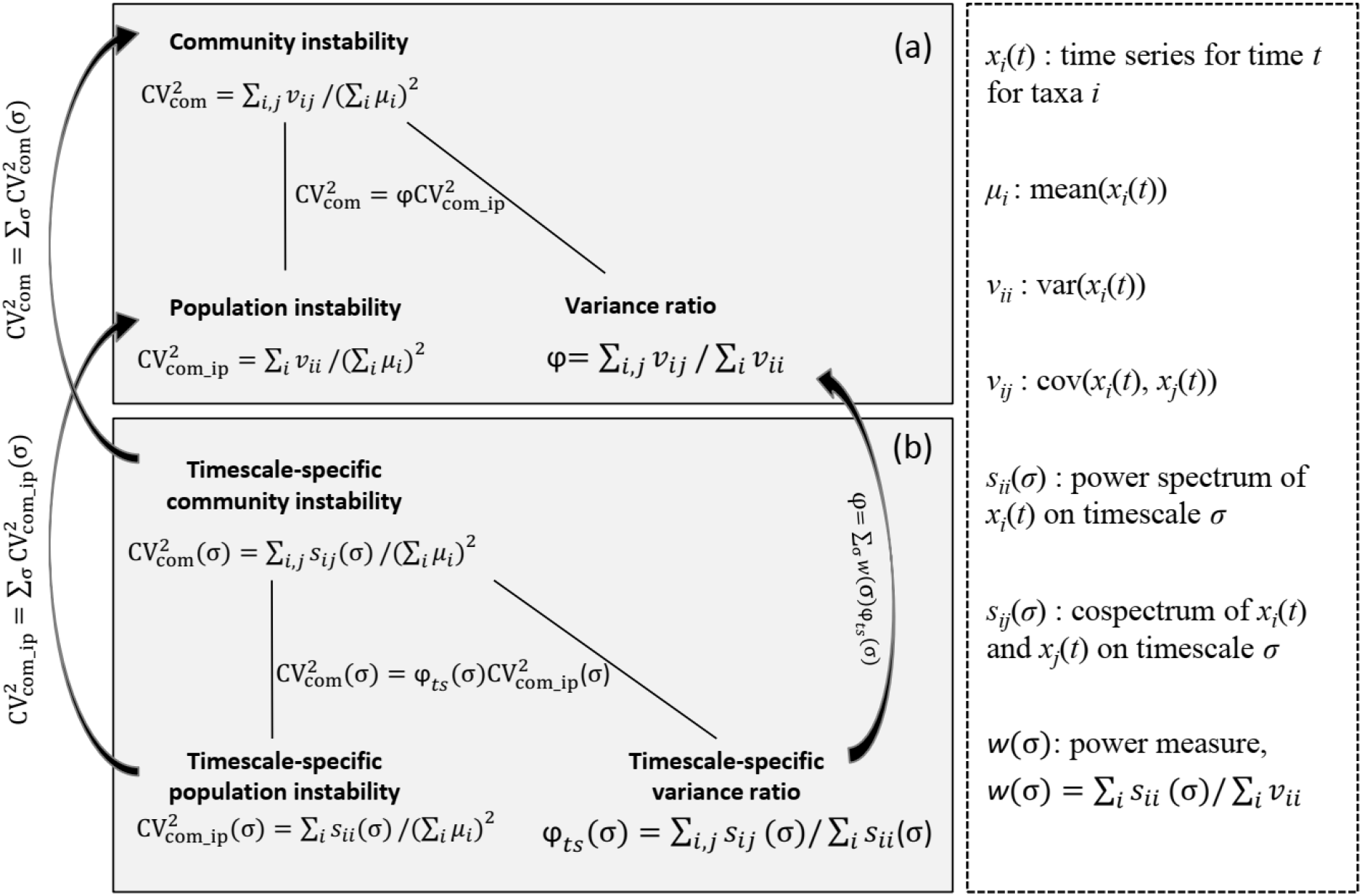
Expressions of original (a) and timescale-specific (b) community variability, aggregate population variability and the variance ratio, as well as the relationships between these quantities.

Community variability, aggregate population variability, and variance ratio concepts can also be defined for any range or set, Ω, of timescales: 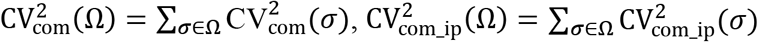, and 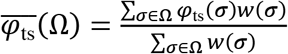. It can then be shown (Appendix S2) that

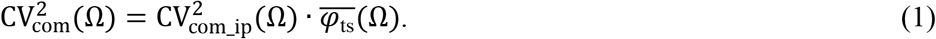

Given a threshold timescale σ_TH_, we define Ω_s_ = {σ: σ < σ_TH_} as “short timescales” and Ω_L_ = {σ: σ ≥ σ_TH_} as “long timescales” relative to the threshold. Equation (1) for Ω = Ω_L_ and Ω = Ω_s_ can then be used to compare the effects of synchrony or compensatory dynamics on population and community variability at long versus short timescales. We note that 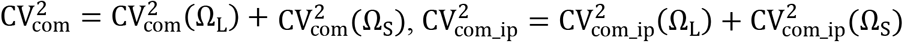, and 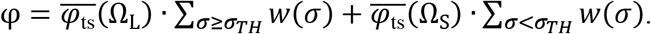.

To illustrate, we apply the theory to the artificial example of Fig. 1. The timescale-specific variance ratio *φ*_st_(*σ*) was greater than one for *σ* = 3, and less than one for *σ* = 10, capturing the deliberately constructed synchronous (*σ* = 3) and compensatory (*σ* = 10) processes at these timescales (Fig. 3a). The non-timescale-specific variance ratio *φ* = 1.064, being a weighted average of *φ_ts_* (*σ*) over timescales, conflated the distinct processes and thereby suggested neither synchronous nor compensatory dynamics. 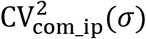 was relatively small for and relatively large for *σ* = 10 (Fig. 3b), reflecting the use of small- and large-magnitude sinusoids, respectively, on these timescales (Fig. 1b,c). The *σ* = 3 oscillations had small magnitude, but were synchronous, whereas the *σ* = 10 oscillations had large magnitude, but were compensatory, hence 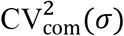 had similar values for *σ* = 3 and *σ* = 10 (Fig. 3c). Community variability at the two timescales was due to weak synchronous oscillations for *σ* = 3 and strong asynchronous oscillations for *σ* = 10 (Fig. 3d); the distinct origins of community variability were revealed by the timescale-specific analysis but not by the classic approach.

**Figure 3.**
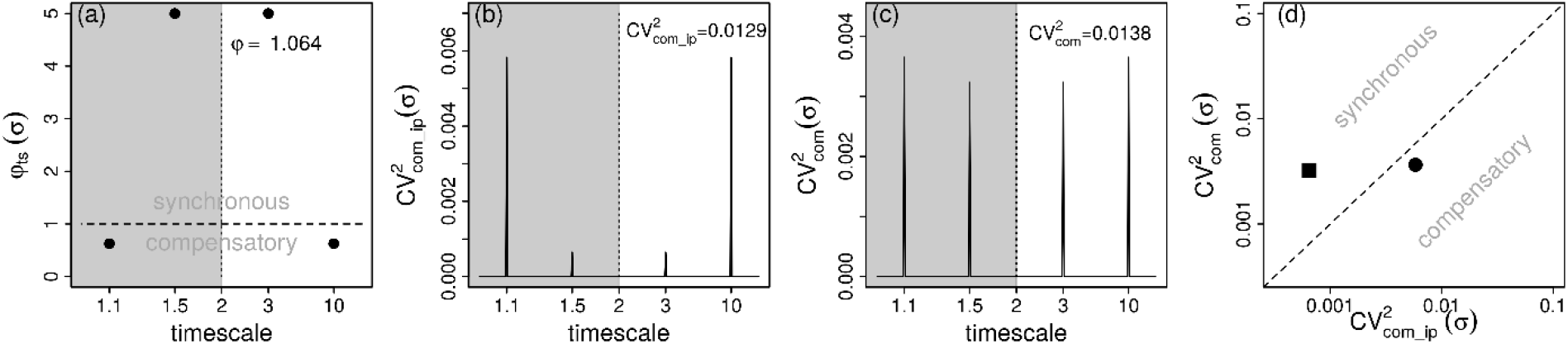
Application of our theory to the artificial time series of Fig. 1. See text of the Theory section for interpretations. Panels a-c show the timescale-specific variance ratio *φ*_ts_(*σ*), aggregate 2 2 population variability CV^2^_com_ip_(*σ*), and community variability CV^2^_com_(*σ*). Timescales less than the Nyquist timescale (gray background) are symmetric to timescales greater than the Nyquist timescale (white background); we present both to reflect the underlying computation of non-timescale-specific quantities (text near top of a-c) as sums or averages across all timescales, both above and below the Nyquist timescale (Theory); but conceptually we focus on the timescales greater than the Nyquist timescale. The horizontal dashed line (a) indicates the boundary between synchronous and compensatory dynamics for each timescale. Panel d compares CV^2^_com_(*σ*) to CV^2^_com_ip_(*σ*) for the period-3 (square) and period-10 (circle) oscillations; the 1:1 line separates synchronous from compensatory dynamics. For simplicity, the example used sums of sinusoids, which oscillate only at discrete timescales, explaining why plotted quantities are 0 (CV^2^_com_ip_(*σ*), CV^2^_com_(*σ*)) or undefined (*φ*_ts_(*σ*)) except at the timescales of oscillation. Real ecological signals are broadband and should typically yield plots that are well-defined and nonzero at all timescales.

## Methods

We applied our timescale-specific variance ratio to six long-term grassland plant community composition datasets from sites distributed throughout the US (Table S1; see Hallett *et al.*, 2014 for detailed descriptions of datasets). Plant abundances were measured either as biomass or percent cover. In cases where abundances were measured as percent cover, summed values were allowed to exceed 100% due to vertically overlapping canopies. All sites were sampled annually (minimum 11 consecutive years) and were spatially replicated (at least 13 plots/site). For sites carrying out longterm experiments, we only used data from unmanipulated control plots. For all sites, methods for data collection were constant over time. One plot was omitted from the Jornada Basin Long Term Ecological Research (LTER) site, because it was an extreme outlier in community variance, leaving 47 plots from that site for our analyses.

For each plot in each site, we calculated the classic variance ratio, *φ*; community variability, 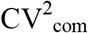; and aggregate population variability, 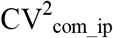. This initial part of our analysis is a replication of part of the analysis of Hallett *et al.* (2014). We also computed our new, timescale-specific measures: the timescale-specific variance ratio, *φ*_ts_(*σ*); the power measure, *w*(*σ*); timescale-specific community variability, 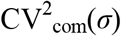; timescale-specific aggregate population variability, 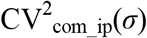; and the band-aggregated quantities 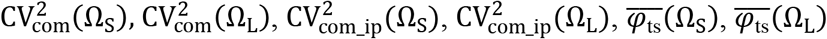. Sheppard *et al.* (2016) used the threshold *σ*_TH_ = 4 years between long and short timescales because that threshold corresponds to a frequency (1 cycle every four years) of exactly half the Nyquist frequency (the maximum rate at which periodic components of the signal can be assessed, which in our case is 1 cycle every two years for annual data), and because it is the boundary between persistent and anti-persistent dynamics (successive values are more similar or dissimilar, respectively) in Fourier components, measured with lag-1 autocorrelation. Here, in applying this theory to the grassland datasets, we use the same threshold. The Fourier quantities *s_ij_*(*σ*) that underlie all of our timescale-specific measures, when not smoothed or averaged across timescales, are highly variable, i.e., plots of these quantities against timescale are “spiky” due to sampling variation. Averaging or smoothing across timescales is the standard statistical approach for ameliorating this property of the *s_ij_*(*σ*), and averaging over wider timescale bands is necessary when data time series are shorter (Brillinger, 2001). We therefore focus our interpretations of results on comparisons between the long and short timescale bands we have defined.

To test whether short or long timescales tended to contribute more to aggregate population variability, we compared the average of 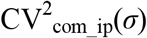 across short timescales (< 4 years) and long timescales (≥ 4 years) for each plot at each site. These quantities are conceptually similar to 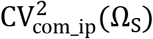, and 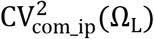 but are averages of 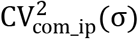 across short and long timescales, respectively, instead of sums. This facilitates comparisons between these quantities: direct comparisons between 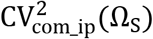 and 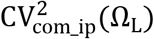 would be complicated by the fact that the sets and can have different numbers of timescales in them, in a way that depends on data time series length. Comparing the average quantities for a single plot indicated whether short or long timescales contributed more, per timescale, to aggregate population variability for that plot. Across all the plots in a site, we conducted a paired t-test to test the significance of the difference between timescales for the site as a whole. If the plots within a site can be regarded as independent replicates, p-value results of these tests have the usual probabilistic interpretation. If spatial autocorrelation or another factor means that plots within a site cannot be regarded as independent, then these p-value results should be interpreted as descriptive statistics, describing the strength of the difference between two paired distributions of values relative to the variation within the distributions. As such, we will refer to these as “nominal” *p*-values.

We used a similar approach to test whether short or long timescales tended to contribute more to community variability. Here we compared the averages of 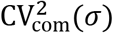 across short and long timescales; these quantities represent the average contributions of a short or long timescale to community variability. Paired *t*-tests were again used to produce nominal *p*-values.

To test whether the degree of synchrony or compensation among populations differed by timescale, we compared 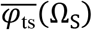 and 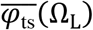. We again used paired *t*-tests and produced nominal *p*-values. These quantities reflect the extent to which either the average or the total, across the timescale band, of aggregate population variability, 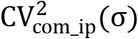, explains the average or the total across the band of community variability, 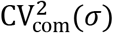. All computations were done in R, all methods are coded into a new R package called tsvr, available at www.github.come/reumandc/tsvr (development version) and on the Comprehensive R Archive Network (CRAN). The package includes a “vignette” that provides step-by-step introduction to use of the package codes. Data are archived at https://portal.edirepository.org/nis/mapbrowse?scope=edi&identifier=358. Codes which can reproduce the results of this study in their entirety at the click of a button are archived at www.github.com/leiku/varrat_decomp. These materials together provide complete transparency and reproducibility for the project.

## Results

### Overall patterns of variability without yet accounting for timescales

Most plots exhibited compensatory dynamics when examined in aggregate across all timescales, although the strength of compensation varied across sites (Hallett *et al.*, 2014). Community variability CV^2^_com_ varied widely, with a minimum observed plot-level value of 0.01 and a maximum of 1.50. Population variability CV^2^_com_ip_ exhibited a similar magnitude of variability, ranging from 0.04 to 1.51. The classic variance ratios *φ* varied from highly compensatory to highly synchronous, with values ranging from 0.08 to 1.98. However, *φ* was less than one in 72.7% of the 150 plots, seemingly indicating (aggregating across timescales) that most communities were less variable through time than would be expected if species fluctuated independently (but see below for a timescale-specific view).

### Timescale-specific patterns of variability

Many plots exhibited marked differences in variability and variance ratios between short and long timescales. To facilitate understanding of results, we first demonstrate this effect with data from one example plot at the Jasper Ridge Biological Preserve (JRG), before later presenting results for all plots. Using the classic approach, dynamics at this plot would be considered compensatory, as the classic variance ratio was 0.457 (which is less than 1), and correspondingly CV^2^_com_ (value 0.047)was less than CV^2^_com_ip_ (value 0.103). However, when we decomposed variability by timescale bands, the short and long timescale bands showed opposite patterns of synchrony and compensation (Fig. 4). The weighted average of *φ*_ts_(*σ*) across short timescales was greater than 1, indicating synchronous dynamics on those timescales, but the weighted average of *φ*_ts_(*σ*) across long timescales was less than 1, indicating compensatory dynamics (Fig. 4a). Correspondingly, the average of CV^2^_com_(σ) was slightly higher than the average of CV^2^_com_ip_(*σ*) when these averages were computed across short timescales, but was substantially less when the averages were computed across long timescales (Fig. 4b-d).

**Figure 4.**
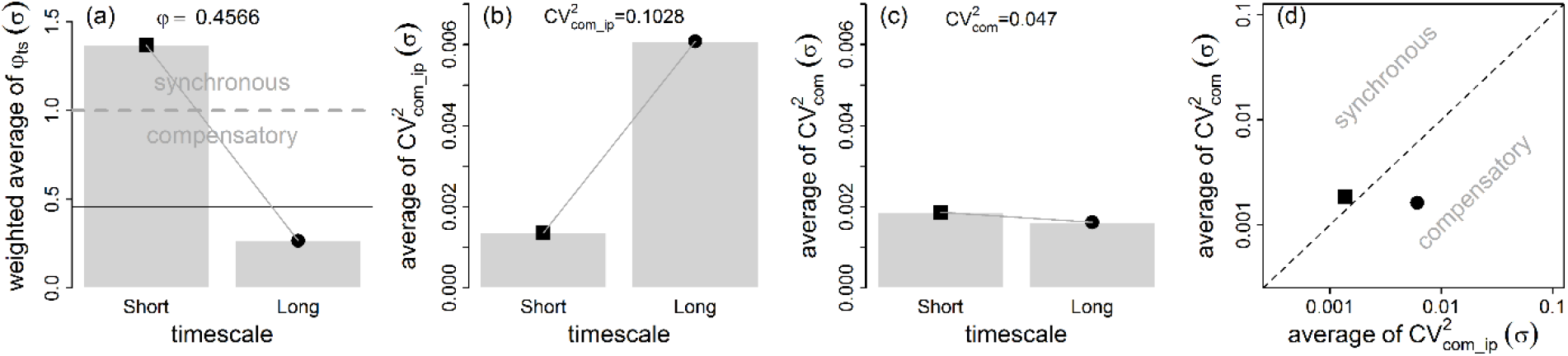
Results for an example plot at the Jasper Ridge Biological Preserve (JRG), which shows a similar pattern to the artificial data of Fig. 1 (see Fig. 3). The classic, non-timescale-specific approach (text at top of panels a-c) suggests compensatory dynamics, and hence CV^2^_com_ is less than CV^2^_com_ip_. But decomposing variability by timescale bands indicated contrasting patterns at short and long timescales: the weighted average of *φ*_ts_(σ) was greater than 1 across short timescales (panel a), so the average of CV^2^_com_(*σ*) (panel c) was slightly greater than the average of CV^2^_com_ip_(*σ*) (panel b) across short timescales; whereas there reverse pattern held for long timescales. Panel d compares the averages of CV^2^_com_(*σ*) and CV^2^_com_ip_(*σ*) across short (square) and long (circle) timescales. See text of the Results section for further interpretation. The horizontal solid line in panel a shows the value of the classic variance ratio.

We applied the same approach to all 150 plots in the six sites (Fig. 5), providing a comprehensive overview of how a timescale approach alters our understanding of compensatory dynamics in these systems. Our data showed average (across plots) greater aggregate population variability at long timescales than at short timescales at all sites (Fig. 5g-l), and likewise for community variability (Fig. 5m-r), except for the Kellogg Biological Station LTER (KBS), where the average CV^2^_com_(*σ*) did not differ between short and long timescales (Fig. 4n). Thus, long timescales were the primary driver of population variability at all sites and of community variability in five of our six sites; this was not surprising given much prior literature on the commonness of temporal autocorrelation in population dynamics (see Discussion).

**Figure 5.**
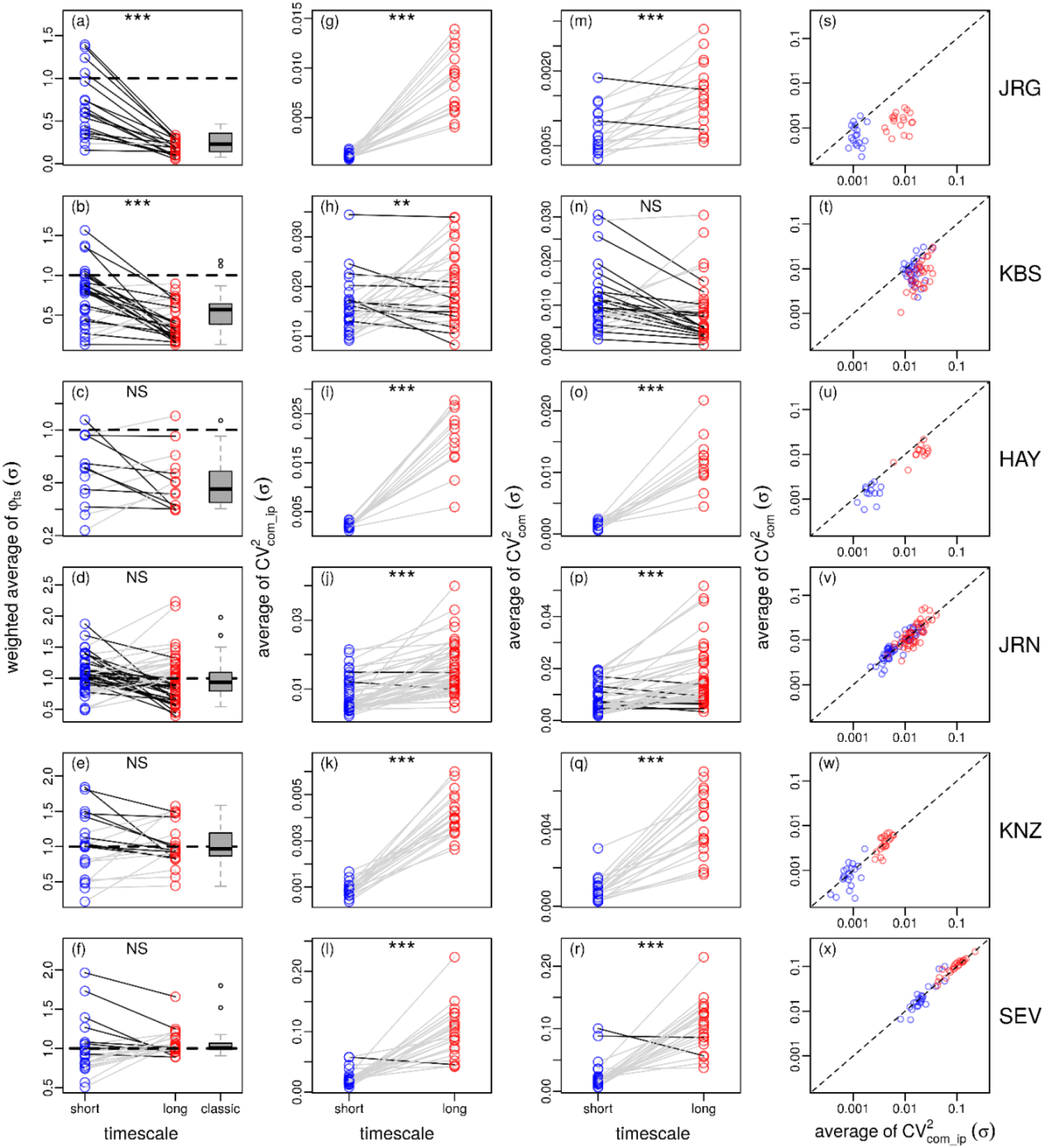
Timescale-specific variance ratio, aggregate population variability and community variability for all plots in each site. Panels parallel those in Figure 4 and show weighted averages of *φ*_ts_(*σ*) (a-f), or averages of CV^2^_com_ip_(*σ*) (g-l) or CV^2^_com_(*σ*) (m-r) across short (blue) or long (red) timescales. Each pair of points on a panel is one plot. Gray lines indicate the value goes up from short to long timescales, while black lines indicate the reverse. Stars in each panel denote nominally significant differences between short or long timescales (paired t-test): *\*\*\*p* <0.001; ** *p* < 0.01; NS: not nominally significant. The box-whisker plots in the first column of panels show distributions across plots of the classic (non-timescale-specific) variance ratio. Panels s-x shows the relationship between the average of CV^2^_com_(*σ*) and the average of CV^2^_com_ip_(*σ*) across short (blue) and long (red) timescales. Site abbreviations: JRG, Jasper Ridge Biological Preserve; KBS, Kellogg Biological Station Long Term Ecological Research (LTER) site; HAY, Hayes, Kansas; JRN, Jornada Basin LTER; KNZ, Konza Prairie LTER; SEV, Sevilletta LTER.

The degree of synchrony and compensation among species differed substantially by timescale at some but not all sites. Short timescales had larger weighted-average values of *φ*_ts_(*σ*) than long timescales for Jasper Ridge (JRG) and Kellogg Biological Station (KBS; Fig. 5a, b; paired t-test, nominal *p* < 0.001), but the weighted averages of *φ*_ts_(*σ*) over short and long timescales were not significantly different at the remaining four sites (Fig. 5c-f; paired t-test, nominal *p* > 0.5). At both JRG and KBS, the weighted average values of *φ*_ts_(*σ*) tended to be < 1 over both short and long timescales, indicating some compensation occurred across all timescales, but compensatory dynamics were much stronger over long timescales.

While Jasper Ridge and Kellogg Biological Station exhibited similar timescale-specific patterns of compensatory dynamics, timescale-specific patterns of community variability differed between these sites for reasons that our approach reveals. At JRG, population variability was much greater on long than on short timescales (Fig. 5g). Community variability was also higher on long timescales (Fig. 5m), but to a lesser degree than population variability, because species dynamics were more compensatory on long than on short timescales (Fig. 5a). At KBS, population variability was also higher on long than on short timescales, but less markedly so than for JRG (Fig. 5h). Because dynamics were again more compensatory on long than on short timescales for KBS (Fig. 5b), community variability was similar on short and long timescales (Fig. 5n). Thus differences across timescales in population variability existed at both JRG and KBS, but opposing differences across timescales in the strength of compensatory dynamics were enough to eliminate timescale differences in community variability for KBS, but only to reduce them relative to population-level differences for JRG.

On average, the remaining four sites (Hayes, HAY; Jornada, JRN; Konza, KNZ; and Sevilleta, SEV) demonstrated similar strengths of compensatory dynamics across timescales (Fig. 5c-f). For these sites, long timescales contributed more to both population and community variability, and to about the same extent (compare Fig. 5i,o for HAY, Fig. 5j,p for JRN, Fig. 5k,q for KNZ, and Fig. 5i,r for SEV). There was substantial variation in weighted-average *φ*_ts_(*σ*) among plots at these sites for both long and short timescales. As a result, there were plots within every site that exhibited a greater tendency toward compensation at long compared to short timescales, and others that showed a lesser such tendency. Full spectral decompositions of all quantities for all plots are given in Figs S1-S6. The distributions, across plots, of all spectral quantities are given for each site in Fig. S7. Comparisons between weighted-average *φ*_ts_(*σ*) and the classic variance ratio *φ* are made for each site in Fig. S8.

## Discussion

A timescale-specific understanding of variability in populations and communities is an important step towards making ecology a predictive science (Kelly and Horton, 2016). Additionally, understanding the links between diversity and stability and the mechanisms that promote diversity and stability are among the long-standing goals of ecology as a field (McNaughton, 1977, Tilman *et al.*, 1998, Ernest and Brown, 2001, Connell and Ghedini, 2015). Although timescale specificity in population dynamics is well-established (e.g., Turchin, 2003, Defriez *et al.*, 2016, Sheppard *et al.*, 2016, Walter *et al.*, 2017), the degree to which community dynamics exhibit timescale specificity is less explored, particularly in terrestrial ecosystems. Timescale specificity in synchrony and compensatory dynamics has been demonstrated in natural and experimental freshwater plankton communities (e.g., Vasseur *et al.*, 2005, Keitt and Fischer, 2006, Vasseur and Gaedke, 2007, Downing *et al.*, 2008, Keitt, 2008, Vasseur *et al.*, 2014, Brown *et al.*, 2016), but methods had not previously been adapted to the shorter, annually sampled time series that are common in terrestrial ecosystems such as grasslands. We developed and applied new methods for the timescale decomposition of the classical variance ratio, using timescale-specific metrics of population and community variability. Although timescale-specific results for six grassland communities show less dramatic differences from results using the classical variance ratio compared to what has been observed in some freshwater systems, our approach did demonstrate the value of considering timescales in 2 of the 6 systems we examined.

In contrast to the methods which have been used to study plankton communities, our methods offer a formal decomposition of the classical variance ratio approach. The classical variance ratio and the community and population variability statistics CV^2^_com_ and CV^2^_com_ip_ can be recovered exactly from our timescale-specific quantities by summing or averaging appropriately over timescales. Thus our approach formally elaborates the classical approach. In developing our theory, we focused on the classical variance ratio (Peterson, 1975, Schluter, 1984). However, a modified variance ratio was proposed by Loreau and de Mazancourt (2008), 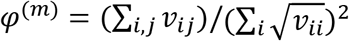, and has become popular in recent years. The alternative approach also uses an alternative to our CV^2^_com_ip_, which we denote 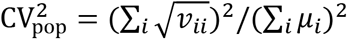 (Loreau and de Mazancourt, 2008). We attempted a timescale-specific extension of the Loreau-de Mazancourt approach to the variance ratio in Appendix S3. Our derivations may facilitate future theory, but are so far not satisfactory for application to data. The fundamental problem is that the CV^2^_pop_ of Loreau and de Mazancourt (2008) does not admit a straightforward timescale decomposition.

Application of our spectral methods to empirical grassland community data showed that plant populations and total community biomass were generally more variable on long timescales than on short timescales. Greater population variability on long timescales is consistent with previous findings that variance increases with time series length in diverse populations (e.g., Pimm and Redfearn, 1988, Halley and Stergiou, 2005). Long time series include variation on longer timescales (Halley, 2005). In natural populations, “red-shifted” (i.e., autocorrelated, or having more spectral power on long timescales) dynamics are common (Steele, 1985, Pimm and Redfearn, 1988, Halley, 1996).

Although some empirical evidence exists for greater community variability on long timescales (Bengtsson *et al.*, 1997), our study appears to be the first observation of this phenomenon in grassland communities. Bengtsson *et al.* (1997) hypothesized that bird community variability was derived from increased population and environmental variability with time, specifically long-term trends in species abundances and temporal heterogeneity in habitat due to succession. In our study, a large proportion of population and community variability was contributed by fluctuations on long timescales (> 4 years). Yet unlike Bengtsson *et al.* (1997), our communities were not successional. Instead, a potential mechanism driving variability on long timescales in our systems may be fluctuations in the environmental resources (e.g., water, nutrients) that affect plant productivity. Resource variation likely propagates through ecosystems, ultimately affecting aggregate community properties in ways that depend on species interactions and shared responses to drivers (Ives, 1995). For example, plant community variability at Jasper Ridge showed a clear spectral peak at a timescale of ~10 years. In contrast, no studies have reported greater community variability at short compared to long timescales. We therefore speculate that community variability may generally be greater at long timescales (>4 years) compared to shorter timescales (< 4 years).

Decomposing community dynamics by timescale may assist in detecting the mechanisms for, and implications of, compensatory dynamics. Compensatory dynamics are promoted by at least two factors: competitive interactions and asynchronous response to environmental change (Ives, 1995, Houlahan *et al.*, 2007, Loreau and Mazancourt, 2013). The latter mechanism is supported by several prior studies in grasslands. For example, Hobbs *et al*. (2007) reported that differential responses of plant species at Jasper Ridge to a period of prolonged below-average rainfall resulted in functional compensation among species. In a broader analysis across nine long-term datasets, Hallett *et al.* (2014) also demonstrated negative covariance among species, especially at sites characterized by high precipitation variability. In systems for which the consequential environmental fluctuations are connected with long-timescale environmental oscillations such as the Pacific decadal oscillation, the timescale signature of compensatory dynamics may help illuminate if the mechanism behind compensatory dynamics is competitive interactions or asynchronous response to environmental fluctuations.

Comparisons between our results for terrestrial grasslands and prior results with freshwater plankton suggest both similarities and differences. First, our finding of stronger compensatory dynamics at some timescales than others is broadly consistent with previous findings in plankton systems that synchrony versus compensatory dynamics depend on timescale (e.g., Keitt and Fischer, 2006, Downing *et al.*, 2008, Vasseur *et al.*, 2014, Brown *et al.*, 2016). Differences by timescale were, however, less profound for grasslands than for the prior plankton studies: no grassland, at the site level, demonstrated a shift from clear compensatory dynamics on one timescale band to clear synchrony on another band, as was observed in both natural plankton populations (e.g., Vasseur *et al.*, 2005) and experimental mesocosms (e.g., Downing *et al.*, 2008). Finally, we did not find strong evidence of synchrony at the site level in any of the systems we studied, unlike the general conclusion that synchronous dynamics prevailed in an analysis of zooplankton in 58 long-term lake datasets (Vasseur *et al.*, 2014).

Our interpretations for grasslands have focused on *average* tendencies within a particular site, either timescale-specificity in the strength of compensatory dynamics (Jasper Ridge, Kellogg Biological Station), or lack thereof (Hayes, Jornada, Konza, and Sevilleta). However, variability across plots within a site was substantial, and every site had some plots that were compensatory on one of the timescale bands we considered and synchronous on the other (Fig. 5). If the variability we observed across plots within a site eclipses sampling variation, then timescale-specific analysis may help reveal, in future work, plot-to-plot heterogeneity in the nature and mechanisms of compensatory dynamics. Exploring potential plot-to-plot variation in the nature of community dynamics probably requires the development of methods to assess the statistical significance of timescale heterogeneities in compensatory dynamics for individual plots. But studying plot-to-plot variation may provide statistical power for illuminating causes of compensatory dynamics that is lacking when making comparisons across whole sites.

Timescale dependency in the presence and magnitude of compensatory dynamics may have implications for how ecologists approach and study ecosystems more generally. For example, synthesis efforts that collate patterns of synchrony and compensation should likely consider length of time series used for such efforts, or should use methods that explicitly take timescale into account. Additionally, do results drawn from short-term observations of communities following manipulation change when observed for longer periods?

To facilitate wider use and adoption of our timescale-specific approach, we developed an open source R package and present this package alongside our paper. Our timescale-specific version of the variance ratio should be particularly attractive for certain ecosystem types and applications. For example, while wavelet methods for quantifying synchrony in ecological communities are increasing in popularity and provide tremendous flexibility (e.g., Vasseur *et al.*, 2005, Keitt and Fischer, 2006, Vasseur and Gaedke, 2007, Vasseur *et al.*, 2014, Brown *et al.*, 2016, Sheppard *et al.*, 2016), these methods are “data hungry”, requiring long time series. Our methods can be used on shorter time series (e.g., >10 equally spaced time points). We hope our results and R package will facilitate researchers of diverse ecosystem types to further illuminate the nature of compensatory versus synchronous dynamics in ecological communities.

## Acknowledgements

This work was part of the LTER Synchrony Synthesis Group funded by the National Science Foundation (NSF) under grant DEB#1545288, through the LTER Network Communications Office, National Center for Ecological Analysis and Synthesis (NCEAS). We thank contributors to the LTER Network; and E. Defriez, J. Dudney, S. Fey, L. Gherardi, N. Lany, M. O’Brien, C. Portales-Reyes, L. Sheppard, and T. Anderson for advice and discussions. D.C.R. and L.Z. were partly supported by the James S. McDonnell Foundation and NSF Grants 1442595 and 1714195. L.Z. was also supported by the Beijing Natural Science Foundation 5194027. K.L.C. was supported by the individual research and development program at NSF. L.G.S. was supported by McDonnell Foundation grant #220020513.

