## Appendix for "A new variance ratio metric to detect the timescale of compensatory dynamics"

#### S1 Basics of Fourier methods

If $x_i(t)$ is a population abundance measurement for taxon $i$ ($i = 1, \ldots, S$) at time $t$ ($t = 1, \ldots, T$), the discrete Fourier transform is

$$X_i(h) = \sum_{t=1}^{T} x_i(t) \exp(-2\pi i(t-1)(h-1)/T), \tag{1}$$

for $h = 1, \ldots, T$. The value $X_i(1)$ equals the sum of the time series $x_i(t)$, and we discard that value. The value $X_i(h)$ for $h > 1$ is the Fourier transform at frequency $(h-1)/T$ (units of cycles per sampling interval), or timescale $\sigma = T/(h-1)$ (units of sampling intervals), which we henceforth denote $X_i(\sigma)$ for $\sigma = T/(T-1), T/(T-2), \ldots, T/2, T$.

We define $s_{ii}(\sigma) = \bar{X}_i(\sigma)X_i(\sigma)/(T(T-1))$, where the overbar represents complex conjugation. If $x_i(t)$ can be assumed to be a finite sample from a stationary stochastic process, then this is the periodogram estimate of the power spectrum of that process [Brillinger, 2001]. We define $s_{ij}(\sigma) = \mathrm{Re}(\bar{X}_i(\sigma)X_j(\sigma))/(T(T-1))$. If the $x_i(t)$ and $x_j(t)$ together can be assumed to be a finite sample from a two-dimensional stationary stochastic process, then this is the periodogram estimate of the cospectrum of that process [Brillinger, 2001].

We now state some facts of basic Fourier analysis without proof. There are many references, including the classic book of Brillinger [2001], that provide proofs and extensive further background on the statistics of spectral analysis of time series. First, $s_{ii}(\sigma)$ and $s_{ij}(\sigma)$ are real-valued quantities, and $\sum_\sigma s_{ii}(\sigma) = \mathrm{var}(x_i(t))$ and $\sum_\sigma s_{ij}(\sigma) = \mathrm{cov}(x_i(t), x_j(t))$, where the summations are over $\sigma = T/(T-1), T/(T-2), \ldots, T/2, T$. Second, $s_{ii}(\sigma)$ and $s_{ij}(\sigma)$ are symmetric about the "Nyquist timescale" (2 time steps), i.e., the vector of those elements of $s_{ii}(\sigma)$ (respectively, $s_{ij}(\sigma)$) for which $\sigma < 2$ equals the vector of those elements of $s_{ii}(\sigma)$ (respectively, $s_{ij}(\sigma)$) for which $\sigma > 2$, written in reverse order. This is why plots of these quantities against timescale are symmetric. Typically one plots and interprets these quantities only for timescales $\sigma \geq 2$, but we plot all timescales in Fig. 3 in the main text because sums and averages over all timescales are required to recover non-timescale-specific variance ratios and population and community variability statistics (see next section), and part of the purpose of that figure is to illustrate that these sums and averages do recover the non-timescale-specific quantities.

#### S2 Details of the theory

We recapitulate definitions from the main text. If $x_i(t)$ is a population abundance measurement for taxon $i$ ($i = 1, \ldots, S$) at time $t$ ($t = 1, \ldots, T$), then let $\mu_i = \mathrm{mean}(x_i(t))$, $v_{ij} = \mathrm{cov}(x_i(t), y_j(t))$, $x_{\mathrm{tot}}(t) = \sum_i x_i(t)$, $\mu_{\mathrm{tot}} = \mathrm{mean}(x_{\mathrm{tot}}(t)) = \sum_i \mu_i$, and $v_{\mathrm{tot}} = \mathrm{var}(x_{\mathrm{tot}}(t)) = \sum_{i,j} v_{ij}$. Then, using the definitions of spectral quantities provided in Appendix S1 and defining $s_{\mathrm{tt}}(\sigma)$ to be the periodogram estimate of the power spectrum of $x_{\mathrm{tot}}(t)$ and

$\Omega$ to be a set of timescales, we define

$$\mathrm{CV}^2_{\mathrm{com}} = \frac{v_{\mathrm{tot}}}{\mu^2_{\mathrm{tot}}} = \frac{\sum_{i,j} v_{ij}}{(\sum_i \mu_i)^2}, \tag{2}$$

$$\mathrm{CV}^2_{\mathrm{com\_ip}} = \frac{\sum_i v_{ii}}{(\sum_i \mu_i)^2} = \frac{\sum_i v_{ii}}{\mu^2_{\mathrm{tot}}}, \tag{3}$$

$$\varphi = \frac{v_{\mathrm{tot}}}{\sum_i v_{ii}} = \frac{\sum_{i,j} v_{ij}}{\sum_i v_{ii}}, \tag{4}$$

$$\mathrm{CV}^2_{\mathrm{com}}(\sigma) = \frac{s_{\mathrm{tt}}(\sigma)}{\mu^2_{\mathrm{tot}}} = \frac{\sum_{i,j} s_{ij}(\sigma)}{(\sum_i \mu_i)^2}, \tag{5}$$

$$\mathrm{CV}^2_{\mathrm{com\_ip}}(\sigma) = \frac{\sum_i s_{ii}(\sigma)}{(\sum_i \mu_i)^2} = \frac{\sum_i s_{ii}(\sigma)}{\mu^2_{\mathrm{tot}}}, \tag{6}$$

$$\varphi_{\mathrm{ts}}(\sigma) = \frac{s_{\mathrm{tt}}(\sigma)}{\sum_i s_{ii}(\sigma)} = \frac{\sum_{i,j} s_{ij}(\sigma)}{\sum_i s_{ii}(\sigma)}, \tag{7}$$

$$w(\sigma) = \frac{\sum_i s_{ii}(\sigma)}{\sum_i v_{ii}}, \tag{8}$$

$$\mathrm{CV}^2_{\mathrm{com}}(\Omega) = \sum_{\sigma\in\Omega} \mathrm{CV}^2_{\mathrm{com}}(\sigma), \tag{9}$$

$$\mathrm{CV}^2_{\mathrm{com\_ip}}(\Omega) = \sum_{\sigma\in\Omega} \mathrm{CV}^2_{\mathrm{com\_ip}}(\sigma), \tag{10}$$

$$\overline{\varphi_{\mathrm{ts}}}(\Omega) = \frac{\sum_{\sigma\in\Omega} \varphi_{\mathrm{ts}}(\sigma) w(\sigma)}{\sum_{\sigma\in\Omega} w(\sigma)}. \tag{11}$$

The quantities $\mu^2_{\mathrm{tot}}$, $\sum_i v_{ii}$ and $\sum_i s_{ii}(\sigma)$ appearing in denominators above should typically be nonzero for all real data sets.

The above definitions lead to three theorems as follows:

**Theorem 1.** *Assuming $\mu_{tot}$ and $\sum_i v_{ii}$ are nonzero, the following timescale decompositions hold exactly, where summations are over the timescales $\sigma = \frac{T}{T-1}, \frac{T}{T-2}, \ldots, \frac{T}{2}, T$:*

$$\sum_\sigma \mathrm{CV}^2_{\mathrm{com}}(\sigma) = \mathrm{CV}^2_{\mathrm{com}}, \tag{12}$$

$$\sum_\sigma \mathrm{CV}^2_{\mathrm{com\_ip}}(\sigma) = \mathrm{CV}^2_{\mathrm{com\_ip}}, \tag{13}$$

$$\sum_\sigma \varphi_{\mathrm{ts}}(\sigma) w(\sigma) = \varphi. \tag{14}$$

*Proof.* Equation (12) follows because

$$\sum_\sigma \mathrm{CV}^2_{\mathrm{com}}(\sigma) = \sum_\sigma \frac{s_{\mathrm{tt}}(\sigma)}{\mu^2_{\mathrm{tot}}} \tag{15}$$

$$= \frac{\sum_\sigma s_{\mathrm{tt}}(\sigma)}{\mu^2_{\mathrm{tot}}} \tag{16}$$

$$= \frac{v_{\mathrm{tot}}}{\mu^2_{\mathrm{tot}}} \tag{17}$$

$$= \mathrm{CV}^2_{\mathrm{com}}, \tag{18}$$

where (17) follows by the properties of the power spectrum estimate (Appendix S1). Equation (13) follows because

$$\sum_\sigma \mathrm{CV}^2_{\mathrm{com\_ip}}(\sigma) = \sum_\sigma \frac{\sum_i s_{ii}(\sigma)}{\mu^2_{\mathrm{tot}}} \tag{19}$$
$$= \frac{\sum_i \sum_\sigma s_{ii}(\sigma)}{\mu^2_{\mathrm{tot}}} \tag{20}$$
$$= \frac{\sum_i v_{ii}}{\mu^2_{\mathrm{tot}}} \tag{21}$$
$$= \mathrm{CV}^2_{\mathrm{com\_ip}}, \tag{22}$$

where (21) again follows by the properties of the power spectrum estimate we use (Appendix S1). Equation (14) follows because

$$\sum_\sigma \varphi_{\mathrm{ts}}(\sigma) w(\sigma) = \sum_\sigma \frac{s_{\mathrm{tt}}(\sigma)}{\sum_i s_{ii}(\sigma)} \frac{\sum_i s_{ii}(\sigma)}{\sum_i v_{ii}} \tag{23}$$
$$= \frac{\sum_\sigma s_{\mathrm{tt}}(\sigma)}{\sum_i v_{ii}} \tag{24}$$
$$= \frac{v_{\mathrm{tot}}}{\sum_i v_{ii}} \tag{25}$$
$$= \varphi, \tag{26}$$

where (25) again follows by the properties of the power spectrum estimate (Appendix S1). There is an assumption used here that $\sum_i s_{ii}(\sigma)$ differs from 0 for all timescales $\sigma$, which should typically be true for real datasets. However, for datasets for which $\sum_i s_{ii}(\sigma)$ is 0 for some $\sigma$ (e.g., Fig. 1 in the main text), the summation in equation 14 can be replaced by a summation over timescales for which $\sum_i s_{ii}(\sigma) \neq 0$ to recover the non-timescale-specific variance ratio (Fig. 3, main text). ■

**Theorem 2.** *Assuming $\mu_{tot}$ and $\sum_i v_{ii}$ are nonzero,*

$$\mathrm{CV}^2_{\mathrm{com}} = \varphi \mathrm{CV}^2_{\mathrm{com\_ip}}. \tag{27}$$

*Again assuming $\mu_{tot}$ and $\sum_i v_{ii}$ are nonzero, and for $\sigma$ such that $\sum_i s_{ii}(\sigma) \neq 0$,*

$$\mathrm{CV}^2_{\mathrm{com}}(\sigma) = \varphi_{\mathrm{ts}}(\sigma) \mathrm{CV}^2_{\mathrm{com\_ip}}(\sigma). \tag{28}$$

*Proof.* Equation (27) holds because

$$\varphi \mathrm{CV}^2_{\mathrm{com\_ip}} = \frac{v_{\mathrm{tot}}}{\sum_i v_{ii}} \frac{\sum_i v_{ii}}{\mu^2_{\mathrm{tot}}} \tag{29}$$
$$= \frac{v_{\mathrm{tot}}}{\mu^2_{\mathrm{tot}}} \tag{30}$$
$$= \mathrm{CV}^2_{\mathrm{com}}. \tag{31}$$

Equation (28) holds because, for $\sigma$ for which $\sum_i s_{ii}(\sigma) \neq 0$, we have

$$\varphi_{\mathrm{ts}}(\sigma) \mathrm{CV}^2_{\mathrm{com\_ip}}(\sigma) = \frac{s_{\mathrm{tt}}(\sigma)}{\sum_i s_{ii}(\sigma)} \frac{\sum_i s_{ii}(\sigma)}{\mu^2_{\mathrm{tot}}} \tag{32}$$
$$= \frac{s_{\mathrm{tt}}(\sigma)}{\mu^2_{\mathrm{tot}}} \tag{33}$$
$$= \mathrm{CV}^2_{\mathrm{com}}(\sigma). \tag{34}$$

■

**Theorem 3.** *Assuming $\mu_{tot}$ and $\sum_i v_{ii}$ are nonzero, and for $\Omega$ only containing $\sigma$ such that $\sum_i s_{ii}(\sigma) \neq 0$,*

$$\mathrm{CV}^2_{\mathrm{com}}(\Omega) = \mathrm{CV}^2_{\mathrm{com\_ip}}(\Omega) \overline{\varphi_{ts}}(\Omega). \tag{35}$$

*Proof.*

$$\mathrm{CV}^2_{\mathrm{com\_ip}}(\Omega)\overline{\varphi_{\mathrm{ts}}}(\Omega) = \left[\sum_{\sigma\in\Omega} \frac{\sum_i s_{ii}(\sigma)}{\mu^2_{\mathrm{tot}}}\right] \frac{\sum_{\sigma\in\Omega} \varphi_{\mathrm{ts}}(\sigma) w(\sigma)}{\sum_{\sigma\in\Omega} w_\sigma} \tag{36}$$

$$= \left[\frac{\sum_{\sigma\in\Omega}\sum_i s_{ii}(\sigma)}{\mu^2_{\mathrm{tot}}}\right] \frac{\sum_{\sigma\in\Omega} \frac{s_{\mathrm{tt}}(\sigma)}{\sum_i s_{ii}(\sigma)} \frac{\sum_i s_{ii}(\sigma)}{\sum_i v_{ii}}}{\sum_{\sigma\in\Omega} \frac{\sum_i s_{ii}(\sigma)}{\sum_i v_{ii}}} \tag{37}$$

$$= \frac{\sum_{\sigma\in\Omega}\sum_i s_{ii}(\sigma)}{\mu^2_{\mathrm{tot}}} \frac{\sum_{\sigma\in\Omega} s_{\mathrm{tt}}(\sigma)}{\sum_i v_{ii}} \frac{\sum_i v_{ii}}{\sum_{\sigma\in\Omega}\sum_i s_{ii}(\sigma)} \tag{38}$$

$$= \frac{\sum_{\sigma\in\Omega} s_{\mathrm{tt}}(\sigma)}{\mu^2_{\mathrm{tot}}} \tag{39}$$

$$= \mathrm{CV}^2_{\mathrm{com}}(\Omega) \tag{40}$$

■

### S3 Connections to the variance ratio of Loreau and de Mazancourt

The variance ratio of Loreau and de Mazancourt is

$$\varphi^{(m)} = \frac{\sum_{i,j} v_{ij}}{(\sum_i \sqrt{v_{ii}})^2}, \tag{41}$$

and takes values between 0 and 1 [Loreau and de Mazancourt, 2008]. This variance ratio can be related to the classic variance ratio via

$$\varphi = \frac{\sum_{i,j} v_{ij}}{\sum_i v_{ii}} \tag{42}$$

$$= \frac{\sum_{i,j} v_{ij}}{(\sum_i \sqrt{v_{ii}})^2} \frac{(\sum_i \sqrt{v_{ii}})^2}{\sum_i v_{ii}} \tag{43}$$

$$= \varphi^{(m)} \frac{(\sum_i \sqrt{v_{ii}})^2}{\sum_i v_{ii}}. \tag{44}$$

The quantity

$$f = \frac{(\sum_i \sqrt{v_{ii}})^2}{\sum_i v_{ii}} \tag{45}$$

takes values between 1 and $S$ (see lemma 4 below), and characterizes the degree to which the variances $v_{ii}$ are heterogeneous. For instance, when these variances are all the same this term is $S$. When there is an index $i$ such that $v_{ii} \gg v_{jj}$ for all $j \neq i$, $f$ is close to 1. The quantity $f$ does not really relate to synchrony, as it does not depend on covariances or correlations between time series in different locations. Thus $\varphi$ has information about synchrony in it, but it has other information as well.

The previous relationship $\mathrm{CV}^2_{\mathrm{com}} = \varphi \mathrm{CV}^2_{\mathrm{com\_ip}}$ becomes

$$\mathrm{CV}^2_{\mathrm{com}} = \varphi^{(m)} f \mathrm{CV}^2_{\mathrm{com\_ip}}. \tag{46}$$

The quantity $\varphi^{(m)}$ is considered an index of synchrony [Loreau and de Mazancourt, 2008], and this new equation makes it clear that not only are synchrony/compensatory dynamics implicated in to what extent population variability ($\mathrm{CV}^2_{\mathrm{com\_ip}}$) translates into community variability ($\mathrm{CV}^2_{\mathrm{com}}$), so is the quantity $f$, which represents the degree to which the variances $v_{ii}$ are heterogeneous. But this makes sense because when these variances are very different, the degree to which time series can reinforce each other (synchrony) or cancel each other out (compensatory dynamics) is limited. Whereas when the variances $v_{ii}$ are similar, time series can reinforce each other or cancel each other out substantially, and $f$ is large and the effects of the overall variance ratio $\varphi$ are accentuated. Following Loreau and de Mazancourt [2008], we define $\mathrm{CV}^2_{\mathrm{pop}} = f\mathrm{CV}^2_{\mathrm{com\_ip}} = \frac{\left(\sum_i \sqrt{v_{ii}}\right)^2}{\left(\sum_i \mu_i\right)^2}$, so that $\mathrm{CV}^2_{\mathrm{com}} = \varphi^{(m)}\mathrm{CV}^2_{\mathrm{pop}}$. Then $\mathrm{CV}^2_{\mathrm{pop}} = \frac{\left(\sum_i \sqrt{v_{ii}}\right)^2}{\left(\sum_i \mu_i\right)^2}$.

The timescale-specific variance ratio can also be decomposed in a similar way,

$$\varphi_{\mathrm{ts}}(\sigma) = \frac{\sum_{i,j} s_{ij}(\sigma)}{(\sum_i \sqrt{s_{ii}(\sigma)})^2} \frac{(\sum_i \sqrt{s_{ii}(\sigma)})^2}{\sum_i s_{ii}(\sigma)}. \tag{47}$$

The first factor on the right here is a timescale-specific version of $\varphi^{(m)}$, which we denote $\varphi^{(m)}_{\mathrm{ts}}(\sigma)$. The new term

$$f(\sigma) = \frac{(\sum_i \sqrt{s_{ii}(\sigma)})^2}{\sum_i s_{ii}(\sigma)} \tag{48}$$

is a timescale-specific version of $f$, with similar interpretation. We can define $\mathrm{CV}^2_{\mathrm{pop}}(\sigma) = f(\sigma)\mathrm{CV}^2_{\mathrm{com\_ip}}(\sigma)$, so that $\mathrm{CV}^2_{\mathrm{com}}(\sigma) = \varphi^{(m)}_{\mathrm{ts}}(\sigma)\mathrm{CV}^2_{\mathrm{pop}}(\sigma)$. Then $\mathrm{CV}^2_{\mathrm{pop}}(\sigma) = \frac{\left(\sum_i \sqrt{s_{ii}(\sigma)}\right)^2}{\left(\sum_i \mu_i\right)^2}$. One would hope that the sum of this quantity across timescales would equal $\mathrm{CV}^2_{\mathrm{pop}}$, but that is not the case because of the square root and the square. For this reason we have been unable to find a satisfactory extension of our timescale-specific approach to the Loreau-de Mazancourt variance ratio.

**Lemma 4.** *The quantity $f$ defined above satisfies $1 \leq f \leq S$.*

*Proof.* It suffices to show $1 \leq \frac{(\sum_{i=1}^S a_i)^2}{\sum_{i=1}^S a_i^2} \leq S$ for positive quantities $a_i$. To see that $1 \leq \frac{(\sum_{i=1}^S a_i)^2}{\sum_{i=1}^S a_i^2}$, note that

$$\left(\sum_i a_i\right)^2 = \sum_{i,j} a_i a_j \tag{49}$$

$$= \sum_i a_i^2 + \sum_{i \neq j} a_i a_j \tag{50}$$

$$\geq \sum_i a_i^2. \tag{51}$$

To show the other inequality, it suffices to show

$$\left(\sum_i a_i\right)^2 \leq S \sum_i a_i^2. \tag{52}$$

Dividing by $S^2$, this is

$$\left(\frac{\sum_i a_i}{S}\right)^2 \leq \frac{\sum_i a_i^2}{S}, \tag{53}$$

i.e., the square of a mean is less than or equal to the mean of squares. But that follows from Jensen's inequality. ∎

### References

D Brillinger. *Time Series: Data Analysis and Theory.* Society for Industrial and Applied Mathematics, Philadelphia, 2001.

M Loreau and C de Mazancourt. Species synchrony and its drivers: neutral and nonneutral community dynamics in fluctuating environments. *American Naturalist*, 172:48–66, 2008.

| Site | Abbr. | Years | Yr. rg. | Plots | Plot size | Richness | Measured | Description |
|---|---|---|---|---|---|---|---|---|
| Jasper Ridge Biological Preserve | JRG | 28 | 1983-2010 | 18 | 0.80 | 25.70 | Percent cover | Serpentine grassland |
| Kellogg Biological Station LTER | KBS | 11 | 1999-2009 | 30 | 1.00 | 34.50 | Biomass | Old field |
| Hays, Kansas | HAY | 30 | 1943-1972 | 13 | 1.00 | 22.20 | Percent cover | Tallgrass prairie |
| Jornada Basin LTER | JRN | 20 | 1989-2008 | 47 | 1.00 | 28.50 | Allometric biomass | Desert grassland |
| Konza Prarie LTER | KNZ | 24 | 1983-2006 | 20 | 10.00 | 37.60 | Percent cover | Annually burned tallgrass prairie |
| Sevilleta LTER | SEV | 13 | 1999-2011 | 22 | 1.00 | 13.40 | Biomass | Desert grassland |

Table S1: Summary of datasets. Plot size in square meters. Richness is the number of species that were ever seen in a plot, averaged across plots for a site. Biomass, when measured, was in g per square meter

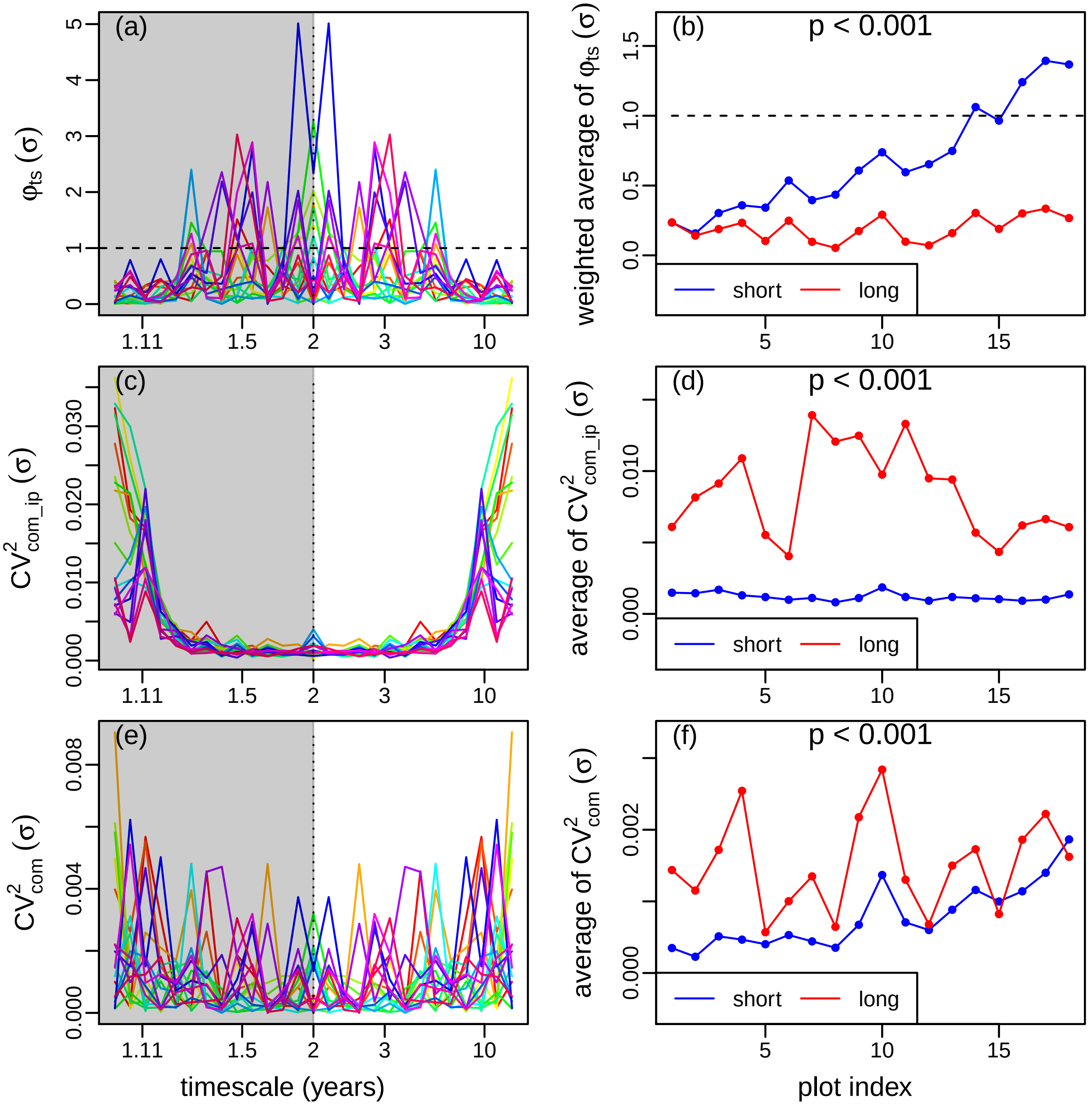


Figure S1: Demonstration of timescale analyses of data from JRG, which consist of 18 plots (indicated by different colors in left panels, and by x-axis in right panels). Different colors in the right panels indicate the average (or weighted average) of the values in the corresponding left panels across short timescales (< 4 years; blue) or long timescales ($\geq$ 4 years; red). On right panels, plots are sorted on the x-axis by the difference between short- and long-timescale averaged $\varphi_{ts}(\sigma)$.

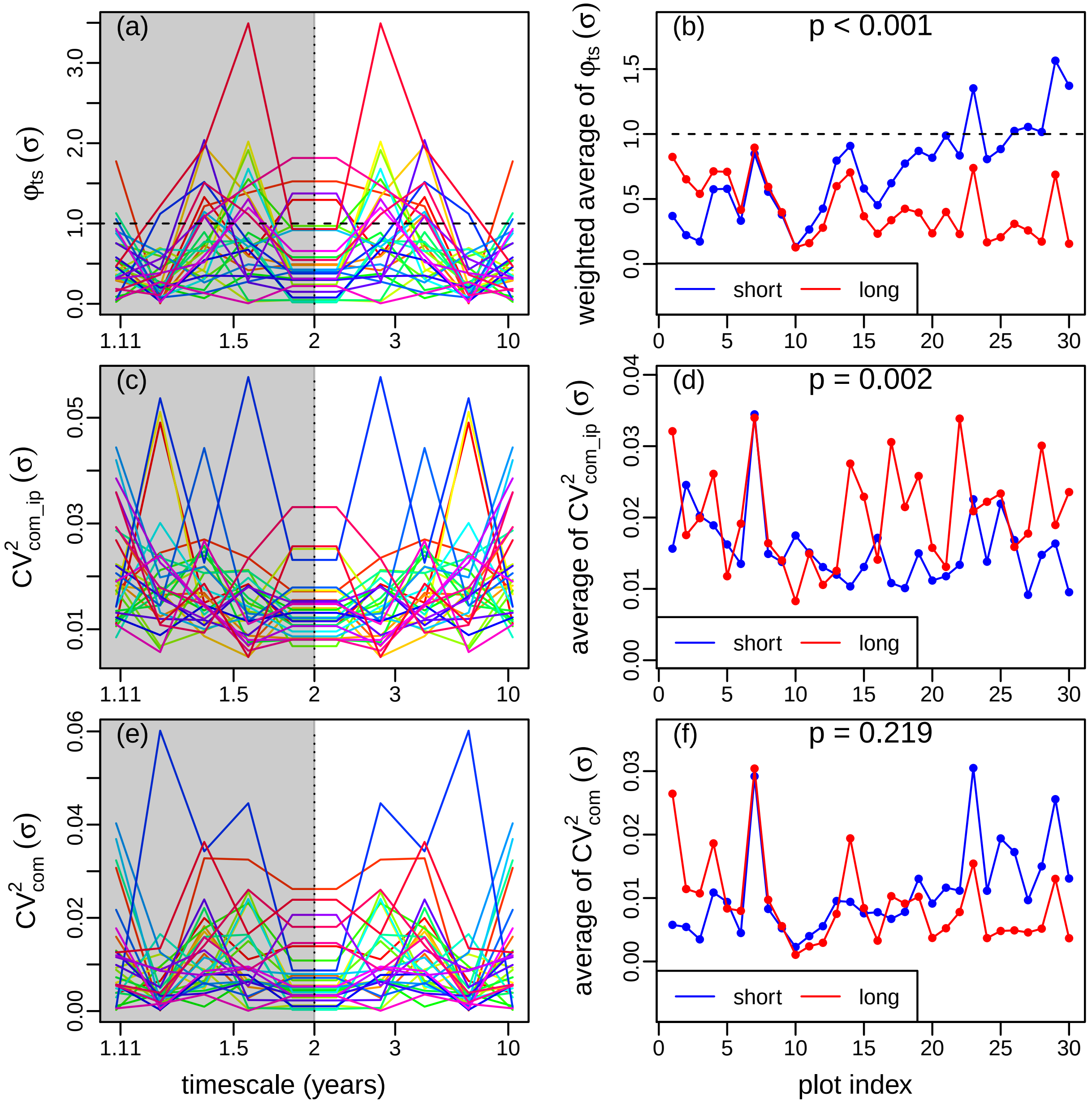


Figure S2: Demonstration of timescale analyses of data from KBS, which consist of 30 plots (indicated by different colors in left panels, and by x-axis in right panels). Different colors in the right panels indicate the average (or weighted average) of the values in the corresponding left panels across short timescales (< 4 years; blue) or long timescales (≥ 4 years; red). On right panels, plots are sorted on the x-axis by the difference between short- and long-timescale averaged $\varphi_{ts}(\sigma)$.

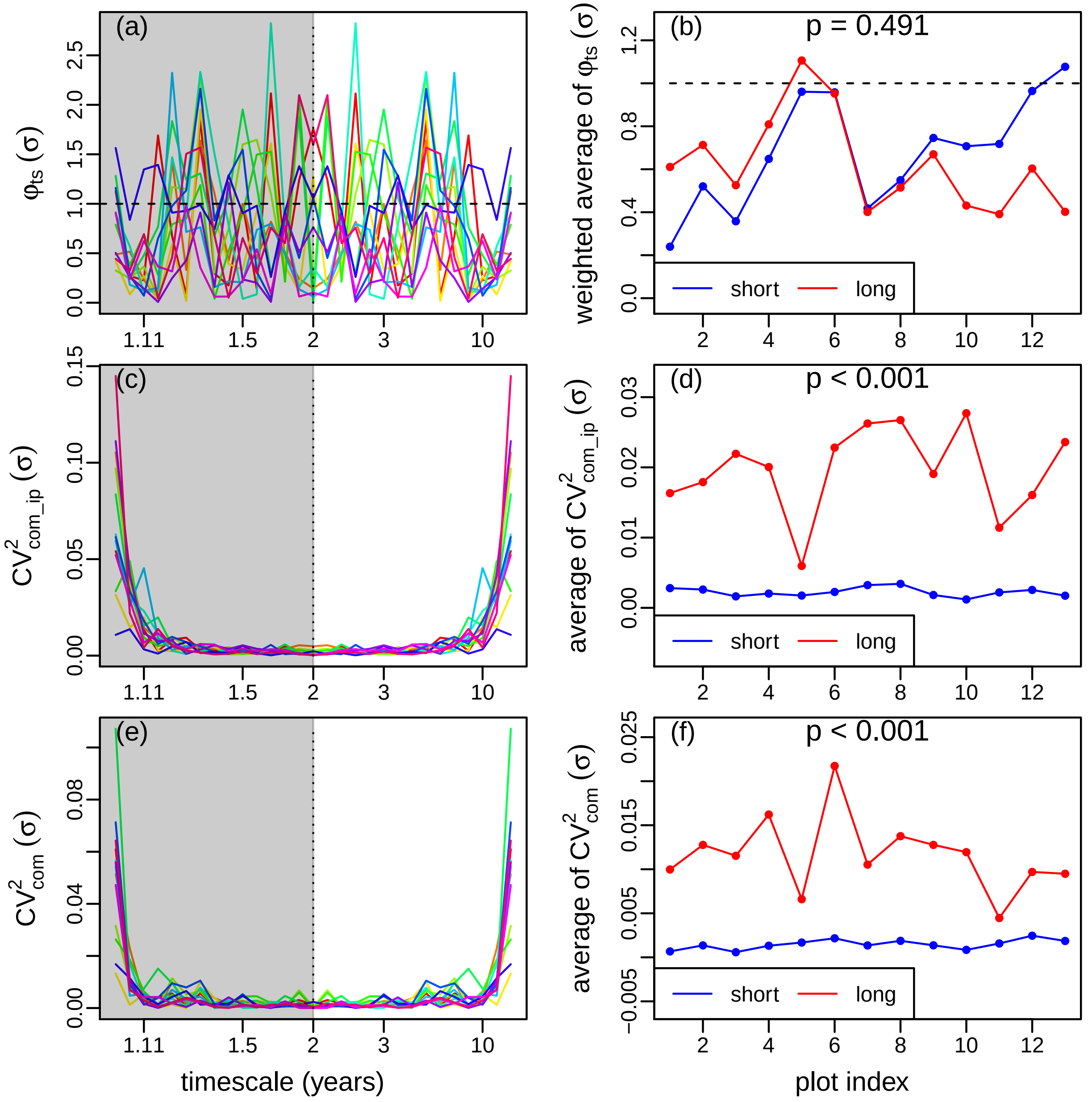


Figure S3: Demonstration of timescale analyses of data from HAY, which consist of 13 plots (indicated by different colors in left panels, and by x-axis in right panels). Different colors in the right panels indicate the average (or weighted average) of the values in the corresponding left panels across short timescales (< 4 years; blue) or long timescales ($\geq$ 4 years; red). On right panels, plots are sorted on the x-axis by the difference between short- and long-timescale averaged $\varphi_{ts}(\sigma)$.

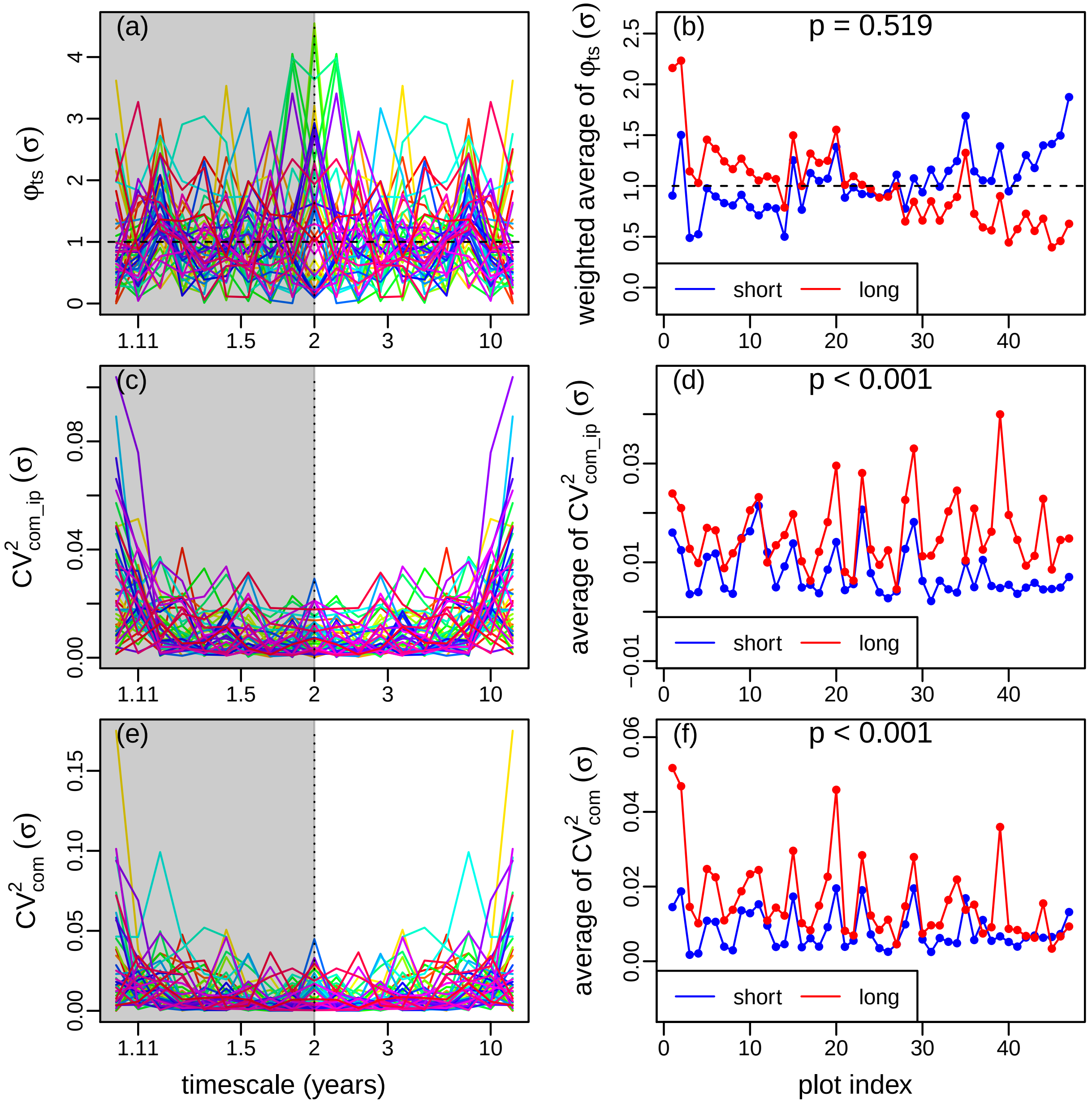


Figure S4: Demonstration of timescale analyses of data from JRN, which consist of 47 plots (indicated by different colors in left panels, and by x-axis in right panels). Different colors in the right panels indicate the average (or weighted average) of the values in the corresponding left panels across short timescales (< 4 years; blue) or long timescales ($\geq$ 4 years; red). On right panels, plots are sorted on the x-axis by the difference between short- and long-timescale averaged $\varphi_{ts}(\sigma)$.

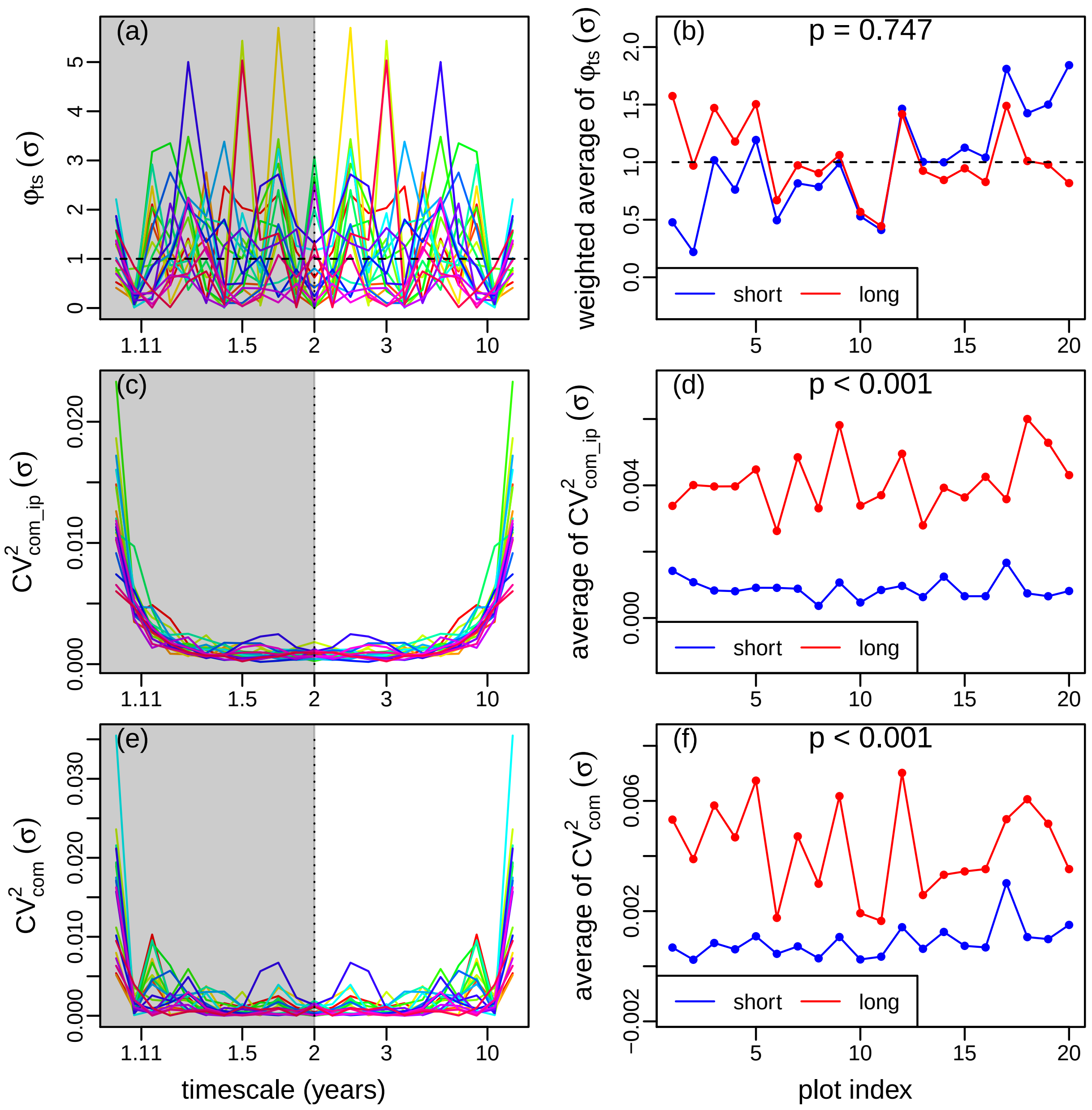


Figure S5: Demonstration of timescale analyses of data from KNZ, which consist of 20 plots (indicated by different colors in left panels, and by x-axis in right panels). Different colors in the right panels indicate the average (or weighted average) of the values in the corresponding left panels across short timescales (< 4 years; blue) or long timescales ($\geq$ 4 years; red). On right panels, plots are sorted on the x-axis by the difference between short- and long-timescale averaged $\varphi_{ts}(\sigma)$.

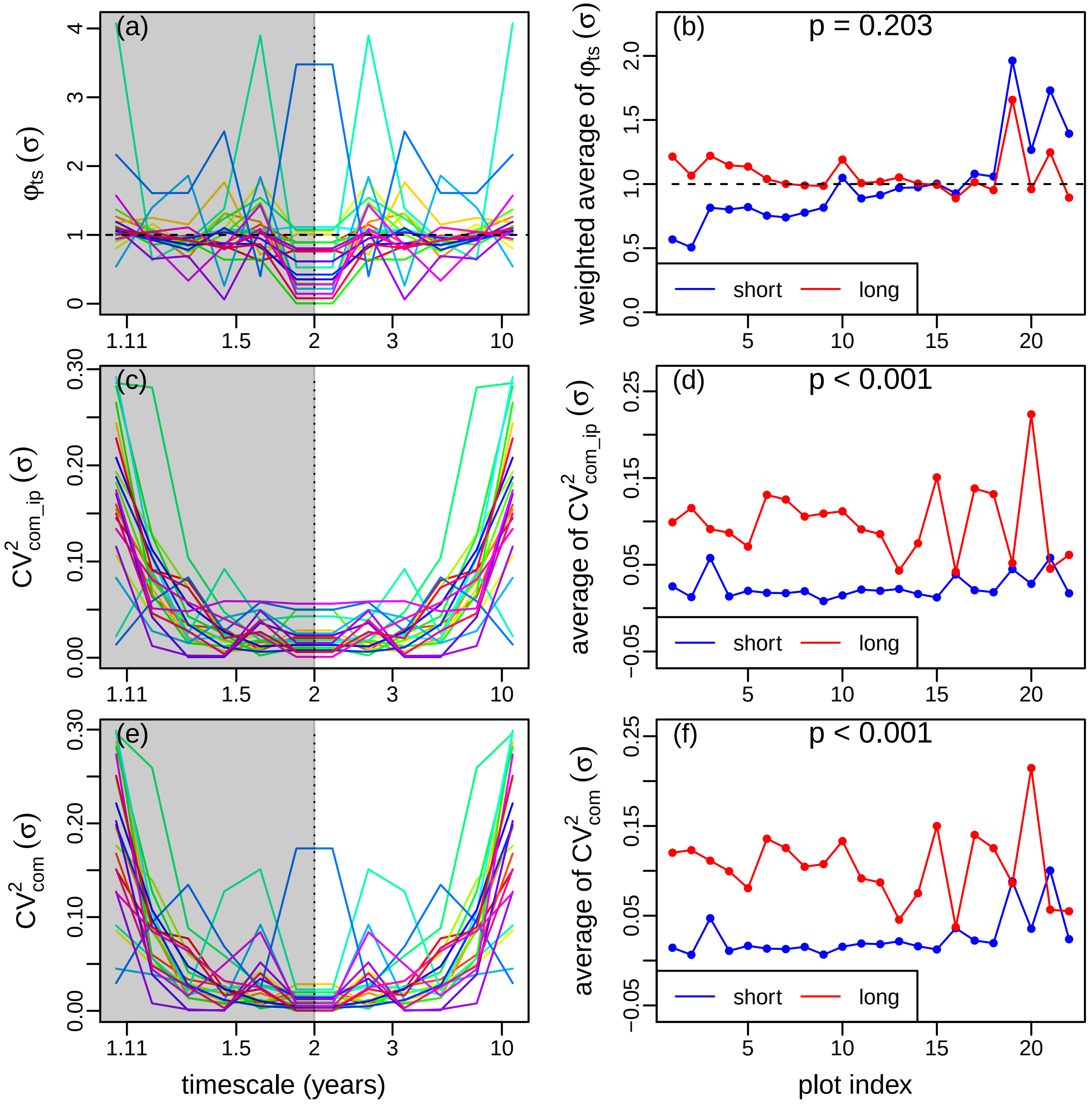


Figure S6: Demonstration of timescale analyses of data from SEV, which consist of 22 plots (indicated by different colors in left panels, and by x-axis in right panels). Different colors in the right panels indicate the average (or weighted average) of the values in the corresponding left panels across short timescales (< 4 years; blue) or long timescales (≥ 4 years; red). On right panels, plots are sorted on the x-axis by the difference between short- and long-timescale averaged $\varphi_{ts}(\sigma)$.

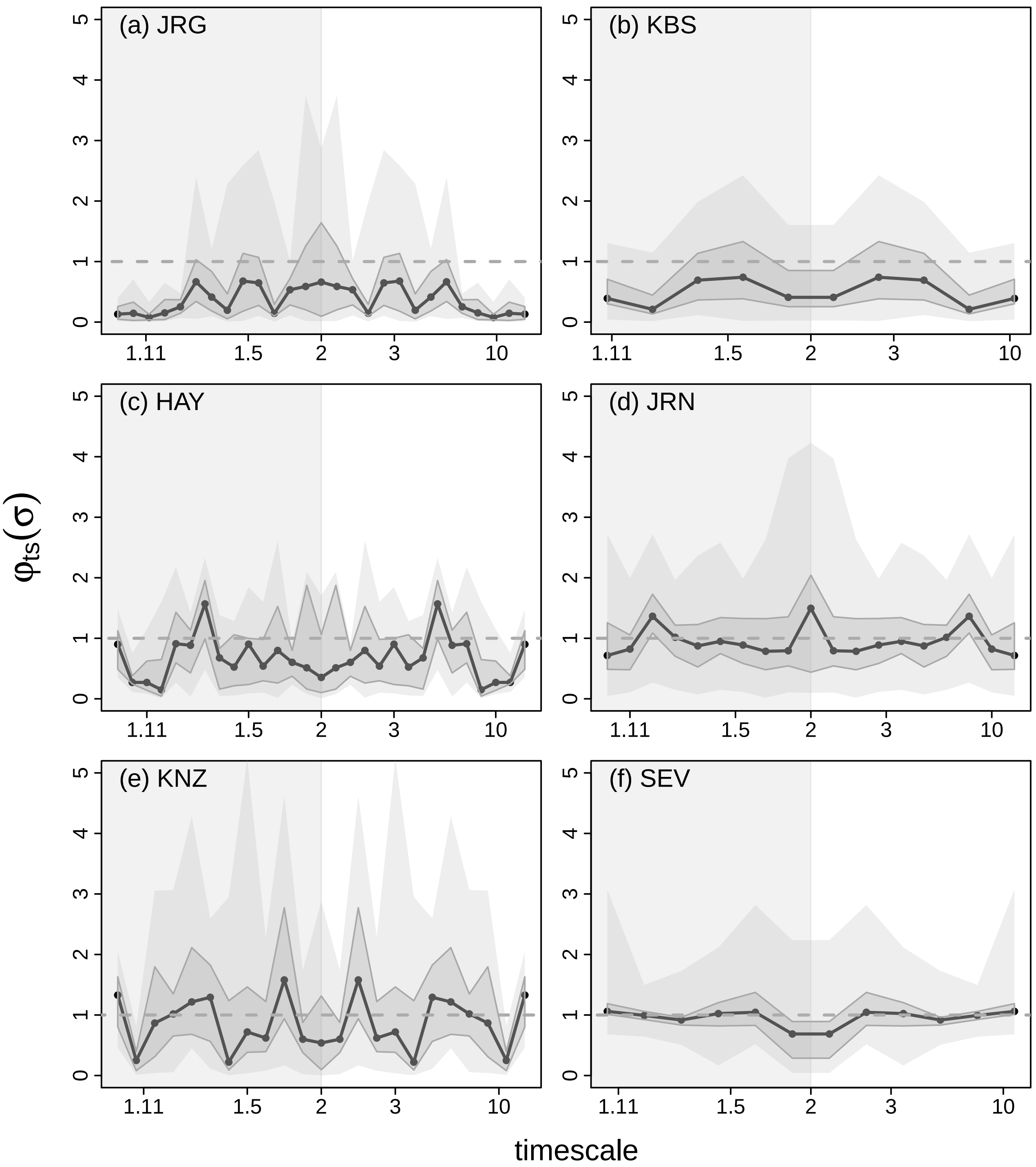


Figure S7: Distributions across plots within sites of timescale-specific variance ratios $\varphi_{\mathrm{ts}}(\sigma)$. The black line is the median of the distribution across plots of $\varphi_{\mathrm{ts}}(\sigma)$ for each timescale, $\sigma$. Dark-gray shading covers $25^{\mathrm{th}}$ and $75^{\mathrm{th}}$ quantiles, and light-gray shading covers $2.5^{\mathrm{th}}$ and $97.5^{\mathrm{th}}$ quantiles. The horizontal dashed line is a variance ratio value of 1.

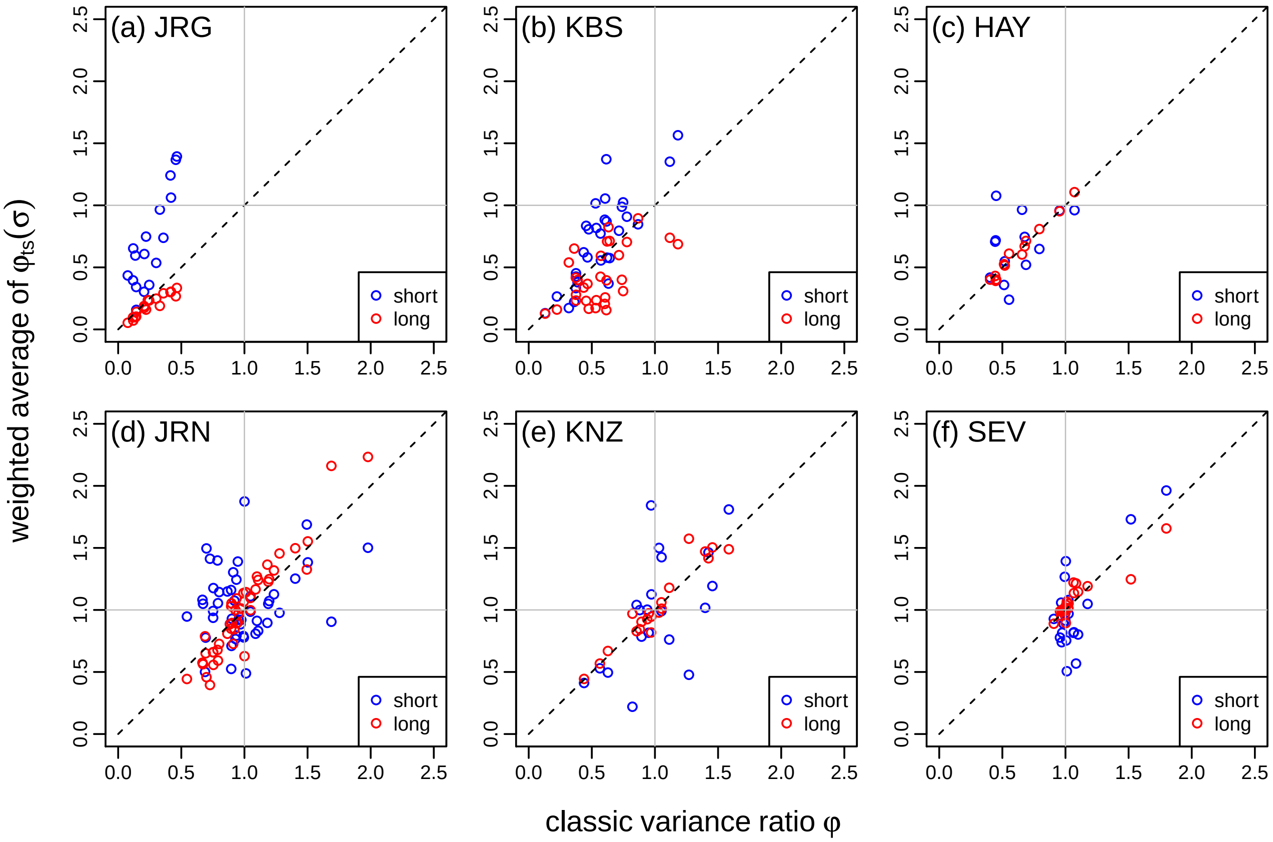


Figure S8: Relationship between the weighted average of $\varphi_{\mathrm{ts}}(\sigma)$ across short (blue) and long (red) timescales and the classic, non-timescale-specific variance ratio across plots for each site. The dashed line on each panel is the 1:1 line.
